# Motivationally objective versus subjective decision-making: Neural correlates, behavior, and self-report

**DOI:** 10.64898/2026.08.22.746476

**Authors:** Priyamvada Modak, Joshua W. Brown

## Abstract

In this study, we investigated the neural and behavioral basis of motivationally objective versus subjective value-based decisions. Using a within-subject fMRI design, healthy participants performed a risky decision-making task that elicited different levels of subjectivity in decision-making in two task conditions. In the ‘Best’ or objective condition, choices were rewarded only when they were objectively best on a given trial, incentivizing decisions based on externally specified per-trial point maximization. In the ‘Choice’ or subjective condition, participants received the reward associated with the chosen option, irrespective of how it compared to the unchosen option, allowing greater freedom to exercise subjective preferences in decision policy. Behaviorally, participants relied more on objectively optimal policy in the Best than the Choice condition. There was also a greater consensus across participants in behaviorally displayed and self-reported policies in the Best condition as well as a greater commitment to a single policy by individual participants in this condition, further confirming more objective behavior in the Best condition, compared to Choice. Moreover, behavioral inferences showed a greater agreement with self-report in the Best condition. Our fMRI results showed that the decision-making in Choice, relative to the Best condition, was associated with greater BOLD response in mid-cingulum/posterior cingulate cortex and dorsal anterior cingulate cortex, suggesting their involvement in less externally constrained, or motivationally subjective, decision-making.

## Introduction

Objectivity is defined as the “Tendency to base judgments and interpretations on external data rather than on subjective factors, such as personal feelings, beliefs, and experiences” (APA). Therefore, we define motivationally objective decisions as those where one relies on ‘facts’ – that is, a valuation accessible to other people and grounded in external or a shared notion of reality. One would be choosing objectively if they were deciding based on principles that they believe are commonly accepted. On the other hand, motivationally subjective decisions stem from a valuation that includes personal preferences and considerations that are unique to oneself and not necessarily shared with others.

Consider the following example. You are walking down the street when a $50 bill falls from your pocket onto the sidewalk. The reality that is shared with other witnesses on the street is that the bill reads $50, losing $50 is undesirable, and that picking it up prevents this loss. From an objective standpoint, you would choose to pick up the bill. However, imagine now that you have severe back ache and would rather avoid bending down. This is your personal reality and preference that only applies to you and that nobody else knows about. From a subjective standpoint, one might choose not to pick up the bill. Of the many decisions one makes every day, some are purely personal and private while some require more objectivity as they may affect others or affect others’ perception of oneself. Accordingly, it is adaptive to be able to control the level of subjectivity we allow in our decisions.

There is relevant previous work on subjective decisions in value-based decision-making which identified neural correlates of distortion from expected value (EV) maximization, quantified as prospect theory parameters (Paulus and Frank 2006; Hsu et al. 2005, 2009; Venkatraman et al. 2009; Rao et al. 2011; Wang et al. 2019). However, this work cannot be used to directly inform the question of motivationally subjective decision-making as this approach conflates subjectivity with deviations from employing a specific policy, EV maximization. The confounding factor here is the knowledge of or the problem-solving ability to find EV maximization as the optimal policy. Misalignment with EV maximization may not necessarily reflect motivationally subjective decisions. Instead, it could also result from someone who is motivationally objective but simply could not find EV maximization as the best policy.

Here we addressed this gap in the literature and targeted the distinction between motivationally objective and subjective directly. We investigated the neural and behavioral correlates of value-based decisions when the evaluation criteria were objective, as incentivized by a more stringent reward structure where participants received a reward only when their chosen option was truly the best in terms of external (experimenter-defined) criteria vs when participants would receive their actual chosen reward outcome, irrespective of whether it was the best as per the above criteria.

Behaviorally, Smith and Krajbich (2021) reported an interaction between the decision condition, option values, and RT in the eye-tracking patterns, where the conditions corresponded to objective (Weight, Size) versus subjective (Liking) evaluation of food items. Although the decision conditions involved evaluating the stimuli on different characteristics, it provides support for motivationally objective versus subjective decisions involving different behavioral strategies.

Relevant neuroimaging work points to several parts of cingulate as well as ventral and dorsal middle prefrontal cortex that may be differentially involved in these conditions. BOLD signals in dorsal anterior cingulate cortex (dorsal ACC) have been reported to correspond to more rational behavior, such as choosing options inconsistent with framing effects (De Martino et al. 2006) or when there is greater linearity of the probability weight function (Kahneman and Tversky 1979; Paulus and Frank 2006). This may suggest a role of dorsal ACC in implementing objective strategies or broadly in selecting choice-strategies or evaluation-criteria. The latter is supported by the observations of Shenhav and Buckner (2014), where greater BOLD signal in dorsal ACC during value-based decisions was associated with a greater likelihood of choice reversals later on. This is also consistent with the findings of Guggisberg et al. (2007) where pregenual and supragenual ACC showed a greater involvement in preferential decision-making, akin to subjective choice, as compared to the ‘numeric’ condition which was akin to an objective choice. Research on subjective valuation of options in value-based decisions points to posterior cingulate cortex (PCC), specifically in subjective valuation of delayed rewards (Kable and Glimcher 2007; Sripada et al. 2011), in signaling integrated subjective value of multi-attribute options (Park et al. 2011), and in signaling learned values from personally experienced rewards over values explicitly shown on the screen (Fitzgerald et al. 2010). The mid-cingulum has also been shown to be involved in emotional awareness (Smith, Ahern, and Lane 2019), along with ACC, implicating these in subjective valuations. Involvement of ACC and PCC in subjective processing is also supported by the research in moral judgement (Hutcherson et al. 2015; Greene et al. 2001). Additionally, previous studies suggest involvement of ventral medial pre-frontal cortex (vmPFC) and orbitofrontal cortex (OFC) in integrating different contributions to the value. Their BOLD activity during decisions has been associated with a greater rationality, that is, smaller framing effects (De Martino et al. 2006) and subjective valuations of odor stimuli (Grabenhorst and Rolls 2009). Lesions of vmPFC and OFC prevent integration of branding information (Koenigs and Tranel 2008) as well as future consequences (Bechara et al. 1994, 1997) in option valuations, and they have been implicated in representing goals (Valentin, Dickinson, and O’Doherty 2007; Zarr and Brown 2023). Further, studies comparing compensatory and non-compensatory decision-making have shown that regions in dorsal medial PFC (dmPFC) track different properties of the options across these conditions (Rao et al. (2011); Rao et al. (2012)), and that regions in dmPFC show the greater BOLD signal during compensatory decisions in participants who predominantly used non-compensatory strategies (Venkatraman et al. 2009). While objectively- vs subjectively-identified strategies may not necessarily translate to the distinctions of compensatory vs non-compensatory or rational vs heuristic, which were the focus of many of the studies discussed above, the regions being tied to these strategies may be involved in a more general role of strategy selection.

To further clarify the nature of objective versus subjective decisions, we asked healthy participants to make two-alternative forced choice risky-decisions in an objective (‘Best’) and a subjective decision-making condition (‘Choice’). To eliminate any confounds, we administered the same task (risky decision-making), same choice options (Gambles), and same reward type (Monetary) to the same person (within-subject design) in both conditions; the only difference between the conditions being what the participants are asked to do via explicit instructions and reward structures, motivating more objective decision-making in the ‘Best’ condition than in the ‘Choice’. We collected fMRI, behavioral, and self-report data from the participants. The ‘Choice’ condition was like the preference-based decisions in most of the previous work discussed above. In our novel design, we compared that with the ‘Best’ condition, aimed at eliciting more objective decision-making with less reliance on subjective preferences. Based on previous work, we predicted that we would find parts of cingulate cotex, dmPFC, or vmPFC/OFC as being involved to a greater extent in subjective or the ‘Choice’ condition.

## Methods

### Participants

All participants (n=32) provided informed consent as overseen by the IU Bloomington IRB. Participants were recruited via posting flyers around the Indiana University campus and around Bloomington. Two of the participants could not see stimuli correctly due to an issue with the projector inside the scanner and hence were excluded from the analysis, effectively making the sample size as 30 participants (18 females). More details on sample demographics can be found in table S1. Further, due to a technical issue, we did not have the scanner start time for some participants. We used a machine learning approach to deduce this information for these participants by training a support vector machine on participant-data where when this information was available, as described in supplementary methods (section 5).

### Task

The task consisted of selecting between two options, a gamble and a sure thing (figure 1B), during the decision phase. The gamble had two or three possible outcomes, and the outcome values (points) and their probabilities were shown to the participants on the screen. During the feedback phase, outcomes of both the options were shown to the participants, irrespective of their choice. The outcome was always and exactly equal to one of the possible gamble outcomes shown during the decision phase. The sure thing was associated with one outcome, which if selected, was obtained by the participant with complete certainty. The gamble outcome was obtained by playing it out. The reward was in terms of points which were later converted to monetary payment as described in supplementary methods (section 4). The side on which gamble appeared, its orientation and the color of both stimuli was selected randomly for each trial.

**Figure 1:**
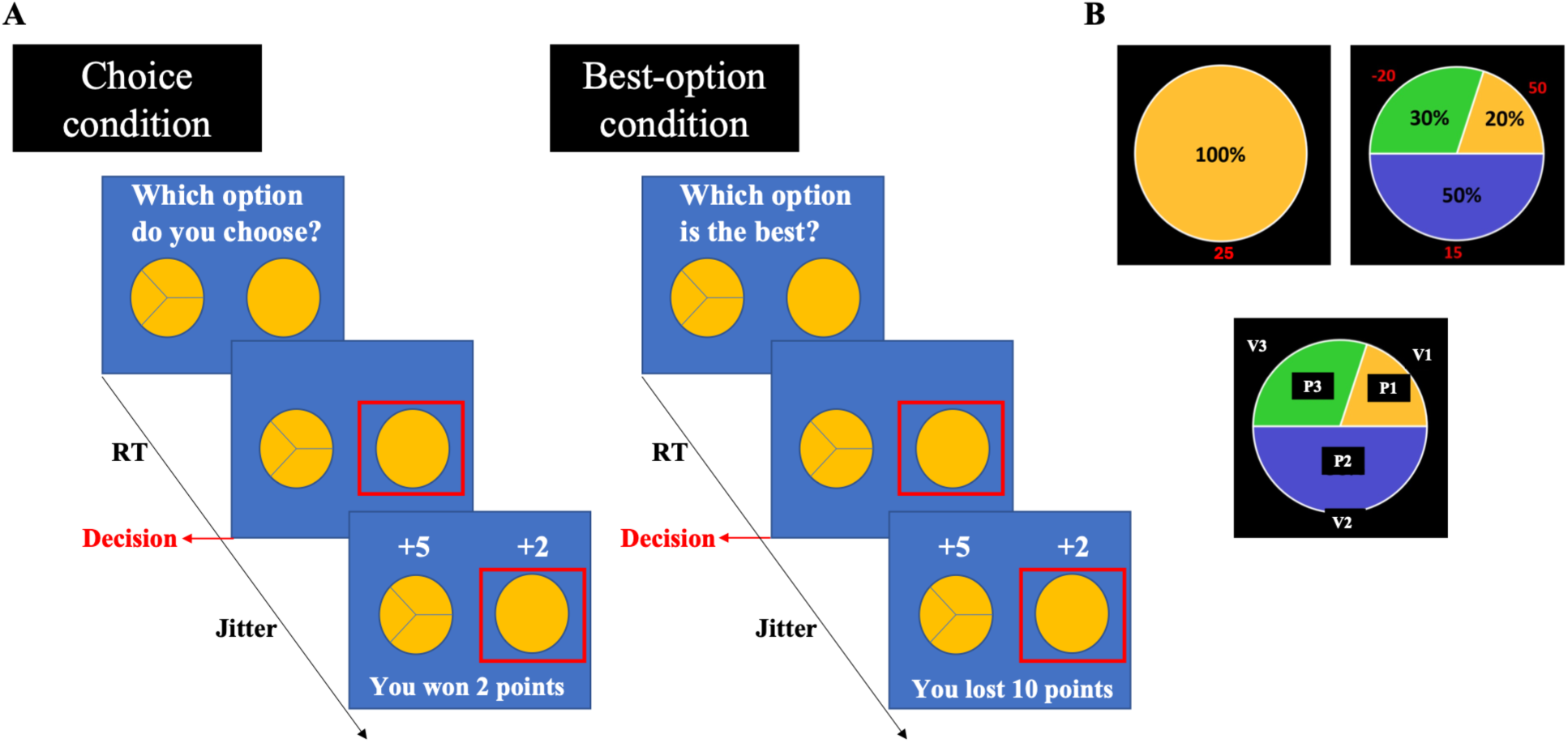
Trial progression and task stimuli. **A.** Sequence of events in a trial of choice and best conditions of our decision-making task. The option with three sections represents the gamble option, and the option with no sections represents the sure-thing option. The values and probabilities associated with each were displayed to the participants on the screen, as shown in B panel. The side (left or right) on which the gamble appeared was selected randomly for each trial. For feedback, the gamble was played out and the outcome of both the gamble and the sure-thing was shown to the participants. **B.** Task stimuli. Top - An example trial showing the gamble (right) and the sure-thing (left). There were five different gambles. Bottom - Different gamble characteristics: V1 - highest positive outcome; V2 - second highest positive outcome; V3 - negative (and the lowest outcome); P1, P2, and P3 are probabilities of V1, V2, and V2 respectively such that P1 + P2 + P3 = 1.

There were three task conditions:

1. ‘Choice’ condition: Participants were asked to choose an option; the outcomes of both the options were shown to them in the feedback phase; the points they earned were same as the outcome of the option they selected, with the gamble outcome determined by sampling from the probabilities as shown. This was adapted from Modak et al. (2021).
2. ‘Best’ condition: Participants were asked to indicate what option is the best; the outcomes of both the options were shown to them in the feedback phase; the points earned depended on how the outcome of the option they selected compared to the outcome of the other option– they received 10 points if the outcome of selected option was more than or equal to the unselected option and lost 10 points otherwise.
3. Control condition: Control blocks showed the same stimuli as the task blocks, but before presenting the stimuli, participants were shown an instruction to select a specific option in the upcoming trial. Control blocks were always conducted after the task blocks and the trials presented here were taken from trials presented in the task blocks. They received 10 points if they selected an option as instructed and lost 10 points otherwise.

Figure 1A displays trial progression in these conditions. The task design aimed at motivating the use of more objective evaluation criteria in the ‘Best’ condition as their choice was penalized if it was not objectively the best between the two options. There was no such penalty in the ‘Choice’ condition, where chose their preferred option and collected the points actually won by the gamble. Participants had 5 seconds to respond.

Each participant completed two consecutive blocks of choice task, two consecutive blocks of best task, and two consecutive blocks of control task, one for choice and one for best. The sequence of choice and best blocks was counterbalanced across participants. Controls blocks were always at the end because the trials presented in the control blocks were randomly sampled from the trials presented during the task blocks and the instructed choice was such that it matched the option participants had chosen in the corresponding task block. This was done to maximize effectiveness of control blocks as controls for stimuli perception and motor action, without any additional effects of their own due to novel options and choice combinations. The blocks of same task condition were conducted consecutively to avoid frequent task-switching and ensure effective experimental manipulation.

There were five specific gamble options, as shown in table S2. For each participant, we estimated two sure-thing values for each gamble using a certainty equivalent task that the participants performed remotely before coming in for the fMRI session, as described below.

#### Certainty equivalent task

Participants completed a certainty equivalent task (Paulus and Frank 2006) online using a personalized link to the webpage we developed for this study. The purpose of conducting this task was to obtain sure-thing values such that the decisions were not too easy for the participants. For each of the five gambles, a participant submitted their choice between the gamble and a sure-thing for a range of sure-thing values. We identified two sure-thing values per gamble corresponding to the choice probabilities 0.35 and 0.65 and used these values with added gaussian noise in the fMRI session. Please see supplementary methods (section 3) for more details.

### Behavioral data analyses

We attempted to discover what decision heuristics participants may have applied. We used each participant’s behavioral data to infer their decision policies during the tasks, from a set of candidate policies (Table S3). Candidate policies included those that represent optimal behavior for our task conditions as well as several heuristics identified in risky decision-making tasks in previous literature (Payne et al. 1992; Brandstätter, Gigerenzer, and Hertwig 2006). The candidate policies were based on: (1) Expected value; (2) Probability of delivering the better outcome (3) Expected gains (4) Expected value of the highest gamble outcome (5) Expected value of the gamble outcomes higher than ST (6) Number of gamble outcomes higher than ST (7) Random choice. Policies 1 and 3-6 had a free parameter ‘c’ representing a measure of risk aversion.

Further, even though participants were explicitly shown the rewards and associated probabilities for both options, it is possible that they were also learning from experience. Therefore, for policies 1–6, learning was incorporated such that the probabilities of the gamble outcomes were learned. As described earlier, participants were shown the outcomes from both options during the feedback phase in all trials, irrespective of their decision. While, in case of gamble, there was uncertainty about what outcome was obtained, there was no uncertainty about the objective value of the specific outcome as they were always exactly same as what was shown in the decision phase. Therefore, learning was incorporated, only for the probabilities of outcomes.

The initial probabilities were assumed to be those explicitly shown to the participants (and which were the ground truth). If the outcome from a gamble was V_i_, the learning happened such that its corresponding probability was increased and the corresponding probabilities of all other outcomes decreased, ensuring that the sum of probabilities of all outcomes remains 1, as shown in equations (1) and (2).

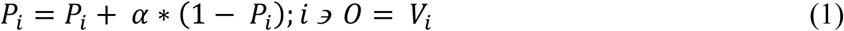

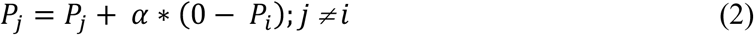

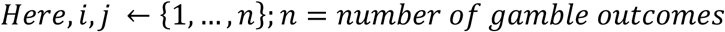

The free parameters for policies 1 and 3 – 6, were c and α, and for policy 2, was α. Both the parameters were assumed to be the same for all gambles for a particular participant.

The optimal parameter values were found such that the log-likelihood of the decisions given the policy was the highest. The parameter space was bounded with α between 0 and 1 and c between 0 and 3. Optimizations were performed at the level of individual participants for both task conditions separately, using genetic algorithm in MATLAB’s global optimization toolbox. Model comparison was performed using Akaike Information Criterion (AIC).

Further, to find the policies that best described the behavior across participants, we pooled together the decisions for all the participants in a particular condition and utilized Bayesian approach to find the policy that was most likely given the data. Bounds on the parameter space were same as that indicated earlier. Prior (Figure 2A) on the policies was such that it penalized complexity according to AIC.

**Figure 2:**
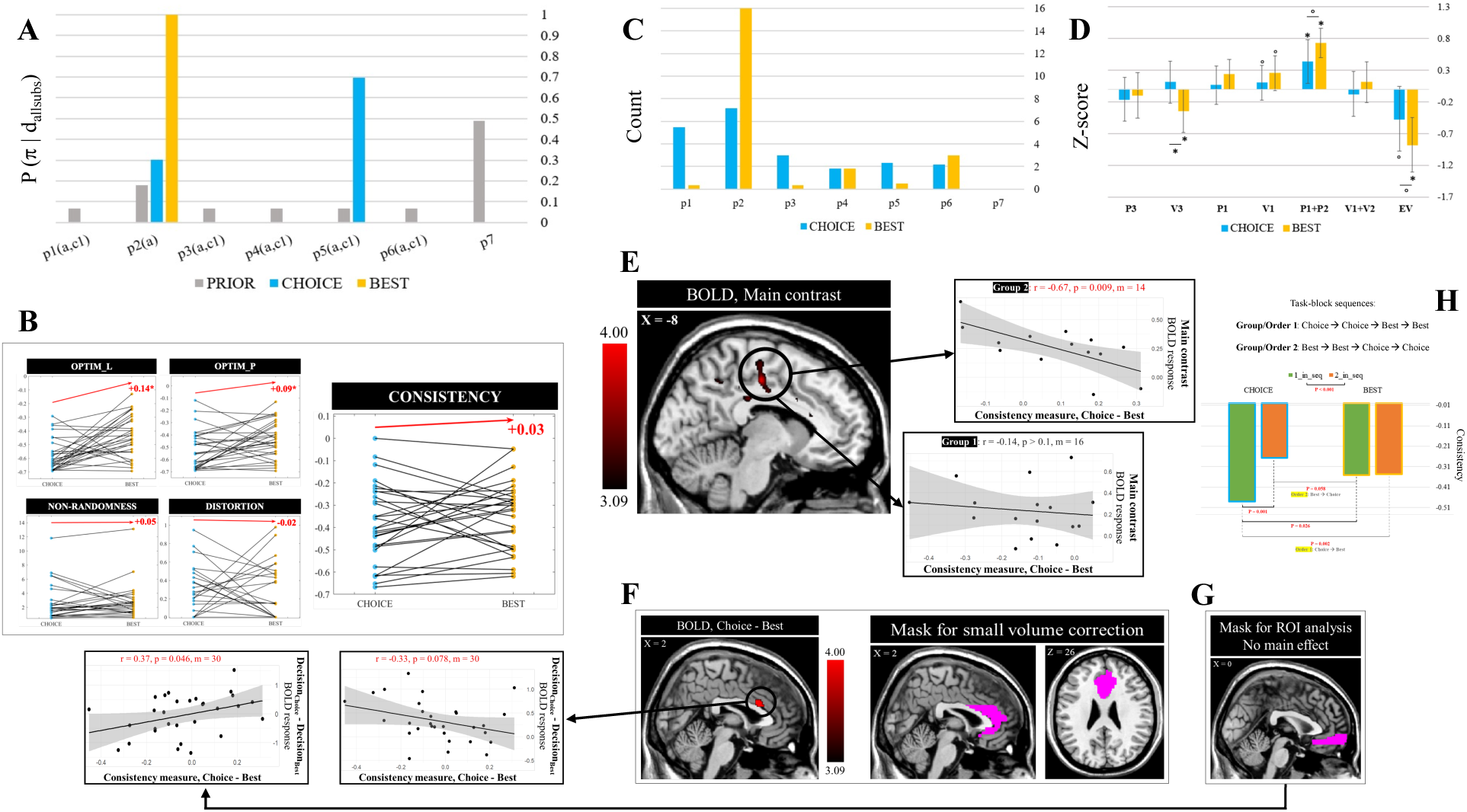
Behavior (A, B, H), self-report (C, D), and neuroimaging (E, F, G) in Choice (subjective) and Best (objective) conditions. **A.** Posterior and prior probabilities of candidate policies for the behavioral data pooled across the participants (supplementary methods, Equation S4). **B.** Objectivity measures obtained from behavioral analysis at the level of individual participants. Red arrow indicates difference between the measures for Choice and Best, averaged across participants. The start of the arrow on y-axis is arbitrary and chosen to be above the data on the plot. ‘*’ indicates significance in the non-parametric paired test at the threshold 0.05. Note: For the non-randomness measure, four participants had extremely high values. They were removed to create the plot. The difference remained non-significant even after including these four participants. **C.** Distribution of policies that the self-reported strategies were mapped to. These were reported by the participants in response to the open-ended questions about task strategies. **D.** Self-reported ratings for the importance of different gamble characteristics in making decisions in the Choice and Best conditions. The ratings corresponding to a particular individual and a particular condition were transformed to z-scores. Here ‘*’ reflects significance at the threshold 0.05 while ‘o’ reflects a trend (p < 0.10). **E.** BOLD effect of the main contrast in a region in mid-cingulate (also extends into posterior cingulate): Region showing significantly positive effect of the main contrast, (DecisionChoice – DecisionChoice_control) – (DecisionBest – DecisionBest_control); Effect of main contrast in this region is negatively and significantly associated with differences in consistency measure between the conditions in group 2. This relation is negative but noisier in group 1. **F.** Small volume correction of the BOLD contrast (DecisionChoice – DecisionBest) using an anterior cingulate cortex (ACC) mask revealed significant effects in the dorsal ACC. Effect of this contrast in dACC was trending negatively with the differences in consistency measure between the conditions. This relation was significant after controlling for RT as mentioned in fMRI results section. **G.** Region of Interest (ROI) analysis for bilateral middle orbito-frontal cortex. Does not show any BOLD effect of the main or the (DecisionChoice – DecisionBest) contrasts but shows positive and significant association with differences in consistency between the conditions. **H.** Order effects in consistency measure: ‘1_in_seq’ – when the task blocks were conducted first in the sequence of blocks during the fMRI session (for Choice, this corresponds to participants of group 1, that is, those who performed the tasks in order 1, Choice ➔ Best, for Best, it corresponds to order 2, Best ➔ Choice. ‘2_in_seq’: when the task blocks were conducted second in the sequence of blocks during the fMRI session (for Choice, it occurs in the case of order 2, and for Best, in the case of order 1).

The details of the optimization at the level of individual participants and across participants can be found in supplementary methods (section 6).

Further, we also compared the choice probabilities in ‘Choice’ and ‘Best’ conditions – for each participant, we found the absolute difference of logit-transformed probability of selecting the sure-thing between these conditions.

### Quantifying objectivity

There are several possible approaches to quantify objectivity at the individual level. One is to assess alignment with rational or optimal policies. In this section, we first describe the optimal policy for each task condition and then describe the measures of objectivity at the individual level.

The policy that is theoretically optimal in a particular task condition depends on whether one is maximizing total/long-term accumulated reward or maximizing the reward in a particular trial. As described earlier, in the Choice task a participant earned points equivalent to the outcome of their decision. In this case, an optimal participant, maximizing accumulated reward would act based on expected value of the options, that is, policy 1 with c = 1. If one is maximizing per-trial reward, then the optimal policy in that case will be to select an option that has the higher probability of yielding a relatively higher reward. This would mean comparing the probability of gamble outcomes that are more than or equal to sure thing with the probability of gamble outcomes less than sure thing, that is, applying policy 2. In the Best task, a participant earned (or lost) fixed (= 10) points if the outcome of the selected option was more (or less) than the outcome of the option not selected. Here, in both cases, when maximizing accumulated reward and maximizing per-trial reward, the optimal policy would be policy 2. This is shown in table 1.

**Table 1:** Theoretically optimal policies in each task condition for long-term and per-trial reward maximization.

| Shorthand | Goal | Choice | Best |
| --- | --- | --- | --- |
| L | Long-term (total) reward maximization | $\pi_1(0,1)$ | $\pi_2(0)$ |
| P | Per trial reward maximization | $\pi_2(0)$ | $\pi_2(0)$ |

\*Parenthesis contains parameter-values for $c$ and $\alpha$ , respectively.

One can define ‘objectivity’ at the level of the individual in several ways, and we propose four approaches here. These are not necessarily mutually exclusive.

#### Behavioral alignment with optimal policy

How much does the theoretically optimal policy in each condition account for the behavior? Policies theoretically optimal with both long-term and per-trial goals were considered. This was quantified as the log-likelihood of the behavioral data for these policies. A higher value may indicate higher objectivity.

#### Consistency measure

Log-likelihood of the behavioral data for the winner policy from the model comparison, described in supplementary methods (section 6). This will be high in cases when a single policy was consistently applied throughout the task, as compared to when a participant switched between multiples policies or showed noisy behavior. A higher value of consistency measure may indicate higher objectivity.

We also defined other objectivity measures, such as ‘distortion’ and ‘non-randomness’ based on optimal parameter values of the winner policy, described in supplementary methods (section 6).

### Self-report measures of task-strategy and experience

After the fMRI session, participants filled out a task-strategy questionnaire, which obtained from them the following: (1) Text responses to open-ended questions about their task-strategy, that is, how they performed the task for each task condition, (2) Rating (1 – 10) on how important each of the gamble values and probabilities were to them in their decision-making (3) Rating (1 – 10) for the difficulty level of the decisions in each task condition (only from 12 participants), (4) Rating (1 – 10) for their familiarity with the concept of expected value. They also took a math aptitude test. The task strategy questionnaire and the math aptitude test can be found in supplementary material.

We mapped participants’ text responses about task-strategy to the candidate policies, and the mapping rule is shown in table S4.

As mentioned above, participants also provided a rating between 1-10 for each gamble characteristic separately to indicate how much it affected their decisions. For a particular participant, their ratings for the gamble characteristics for a condition were z-scored. This provided a way to find, across the participants, which characteristic was consistently reported as the most (or least) important, relative to the other characteristics, in making decisions. The number of participants (n = 27) considered for this analysis were those who had provided ratings for all characteristics in both conditions.

During the fMRI session, we also obtained from 12 participants their rating (1 – 5) for their experienced freedom of choice after each task block. We report on the details and results of this part of the study, in our forthcoming paper (Modak and Brown 2026).

### Agreement between behavior and self-report

We quantified the agreement between self-reported task-strategies and displayed behaviors as follows. First, we computed the log-likelihood of the behavioral data given self-reported policy (at its optimal parameterization), referred as LL_selfreport. This indicated the extent, for a participant and a task condition, to which the self-reported policy accounted for the behavior.

Second, we obtained the posterior probability of the self-reported policy given the behavioral data. In obtaining the posterior probability, the candidate policies at their optimal parameterization were considered (as shown in equation S5), however, with a uniform prior as there was no reason to penalize complexity in measuring agreement between self-report and behavior. This is referred to as posterior_selfreport.

Finally, we determined the rank of the self-reported policy relative to other policies in terms of the log-likelihood the behavioral data for the candidate policies. Note that a higher value of the rank indicates lower agreement. This is referred to as rank_selfreport.

Posterior_selfreport and rank_selfreport indicated, for a participant and a task condition, the extent to which the behavioral data favors the self-reported policy relative to other candidate policies.

The ‘self-reported policy’ above is the policy to which participant’s response to open-ended questions about task-strategy was mapped to.

### fMRI analyses

We performed general linear modeling (GLM) of fMRI data using SPM12 in MATLAB (version 2013a) and created a GLM with ‘Decision’ as an event regressor at the time when participants submitted their decision. We also created additional GLMs where we parametrically modulated the ‘Decision’ regressor with V2, P1*V1, P3*V3, and the value of gamble according to policies 2 and 5. Please see figure 1B for a description of P1, V1, P3, and V2. A separate GLM was constructed and estimated for each parametric modulator.

Our <u>main contrast</u> of interest was (Decision_Choice_ – Decision_Choice_control_) – (Decision_Best_ – Decision_Best_control_).

We also performed small volume correction of the main contrast and (Decision_Choice_ – Decision_Best_) contrast with anterior cingulate cortex and orbito-frontal gyrus masks and ROI analysis with bilateral middle orbito-frontal gyrus.

Details of fMRI acquisition and preprocessing are provided in the supplementary methods (section 9).

## Results

### Behavior

The reaction time (RT) in the Best (objective) condition, across the participants, was significantly more than the Choice condition (Non-parametric paired sample test; Choice – Best, mean = - 151 ms; p = 0.0387). A non-parametric (wilcoxon signed rank) test was performed as the RT was capped at 5 seconds, rendering the normality assumptions of parametric tests invalid. There was no significant difference in choice probabilities between the conditions.

As mentioned in the Methods section, we employed a Bayesian approach to find the policy that best described the behavior across the participants. Figure 2A shows the prior and posterior probability for each of these policies in each condition and corresponds to equation S4 (supplementary methods).

In the Choice condition, policy 5 had the highest posterior probability (P (π5(α,c1) | data_choice_) = 0.70) while in Best, it was policy 2 (P (π2(α) | data_choice_) ∼ 1). We looked at the Bayes’ Factors (BFs) of candidate policies against the winner policy as a measure of how strong the evidence is for the winner over others. We adopted a threshold of 0.1 (Jeffreys 1961; Kass and Raftery 1995), such that when BF was less than 0.1, the evidence for the winner policy was considered strong.

From the BFs (Figure S3), the data in each condition supports the policies in the following order:

**Choice**: π5(α,c1) > π2(α) **>>** π4(α,c1) > π3(α,c1) **>>** π6(α,c1) **>>** π1(α,c1) **>>** π7

**Best**: π2(α) **>>** π5(α,c1) **>>** π6(α,c1) **>>** π4(α,c1) **>>** π3(α,c1) **>>** π1(α,c1) **>>** π7

Here, ‘>>’ represents strong evidence in favor of left over right while ‘>’ represents sub-threshold evidence for favoring the policy on the left.

We also individually fit these policies (Equation S5) to the data pooled across participants, which revealed the optimal parameter values for policy 5 in the choice condition as α = 0 and c1 = 0.78 with (P (π5(0,0.78) | data_choice_) = 0.99; slope_logistic_regression_ = -0.12, p < 0.001). In the Best condition, the optimal parameter for policy 2 was α = 0 with (P(π2(0) | data_best_) ∼ 1; slope_logistic_regression_ = - 3.32, p < 0.001). We also performed additional model comparison at the level of individual participants which allowed inferring the policy and its parameterization that best accounted for the behavioral data for each participant in each condition (Table S6).

There was no difference in learning, inferred from optimal learning rates that were zero for both the conditions for the data across-participants, as reported above. Further, within-sample Wilcoxon signed rank test of the learning rates of the optimal policies estimated at the level of individual participants, across the participants, was also not significant. While there was no significant difference between the conditions, for the Best condition alone the average learning rate was 0.11 (non-parametric, p = 0.0039). There were 8 participants in the Best condition with non-zero learning rate.

### Self-report: Task strategies

As described in the Methods, we obtained participant responses to open-ended questions on task strategy and their ratings of how important different gamble characteristics were in their decision-making in each condition.

In terms of the differences that participants explicitly mentioned between the tasks in their responses, several participants mentioned that they made decisions based more on intuition/gut-feeling/rough-calculations in Choice than in the Best condition, versus being more explorative in Choice. Please see supplementary Table S5 for each participant’s responses to these open-ended questions as well as for condensed take-aways from the responses.

These responses were mapped onto candidate policies, as shown in supplementary table S4.

Figure 2C shows a histogram of the policies that the responses were mapped to, across the participants.

Self-reported ratings for the importance of gamble characteristics were z-scored for a particular subject and a particular condition. This provided a measure of how important a gamble characteristic was relative to other characteristics in a condition for a participant while controlling for inter-subject variability in providing overall ratings. One sample t-test (n = 27) of the transformed ratings for individual gamble characteristics revealed that:

- P1 + P2 had a high importance z-score in both the Choice (p = 0.015) and the Best conditions (p < 0.001), with the difference (Choice < Best) being close to significant (p = 0.078).
- EV had a low importance z-score in both the Choice (close to significant; p = 0.069) and the Best conditions (p < 0.001), with the difference (Choice > Best) being close to significant (p = 0.071).
- V3, or value of loss, had a low importance z-score in the Best condition (p = 0.05), and was significantly lower than the Choice condition (p = 0.012).

This is shown in figure 2D. This indicates that P1 + P2 was consistently rated as important (relative to other characteristics) in both the conditions but its rated importance trended higher in the Best condition than Choice. EV was consistently rated as of low importance in both conditions, but its rated importance trended higher for the Choice condition than the Best. Further, the importance of V3 or the value of loss was rated significantly lower in the Best condition than the Choice condition.

While maximizing expected value is rational in the Choice condition, participants varied in their familiarity with expected value (Figure 3) with several of them reporting to be completely unfamiliar with the concept.

**Figure 3:**
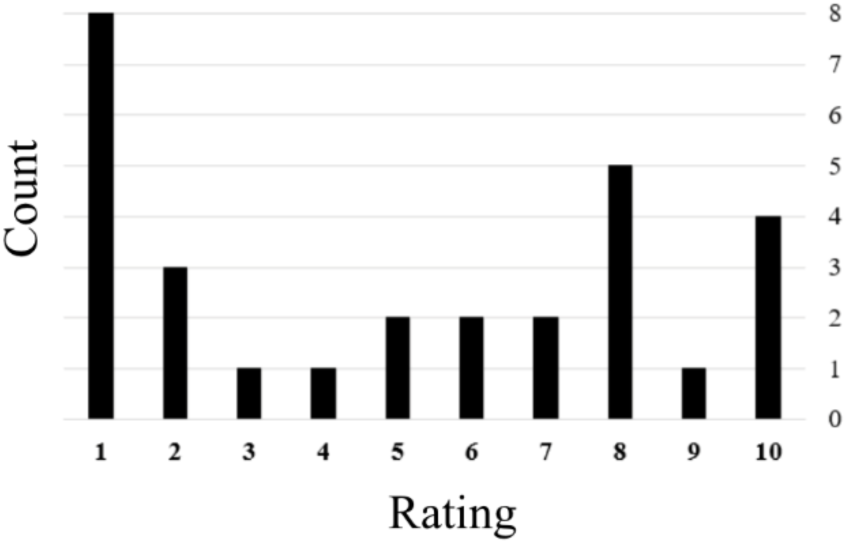
Familiarity with the concept of expected value (EV). 1 – Completely unaware, 10 – Aware and comfortable with calculating it.

There was no significant difference in the self-reported difficulty between the Choice and the Best condition. Difficulty in the Choice condition was significantly greater than control (Choice >Control, mean = 5.75, non-parametric paired test, p < 0.001, n = 12), and in the Best condition too, it was significantly greater than control (Best > Control, mean = 5.83, non-parametric paired test, p < 0.001, n = 12).

### Objectivity measures

We quantified objectivity at the level of individual participants in several ways, as described in Methods. We obtained objectivity measures for each participant individually and compared them between the Choice and Best conditions, as shown in figure 2B. This comparison whne the data is pooled across participants is shown in figure S4. Behavior in the Best condition showed greater consistency, less distortion, less randomness, and greater alignment with theoretically optimal policies for both long-term and short-term goals. Non-parametric paired tests revealed that these differences were significant for the alignment with theoretically optimal policies (non-parametric paired test: Theoret_L, Choice < Best, -0.14, p < 0.001; Theoret_P, Choice < Best, mean = -0.09, p = 0.016).

#### Behavior and self-report

Figure 4 shows that the difference in consistency measure between the Choice and the Best conditions was negatively related to difference in self-reported difficulty between the conditions (r = -0.7, p = 0.01, n = 12) and the RT of the decisions (r = -0.68, p < 0.001, n = 30). Greater consistency was associated with lower difficulty and faster RTs.

**Figure 4:**
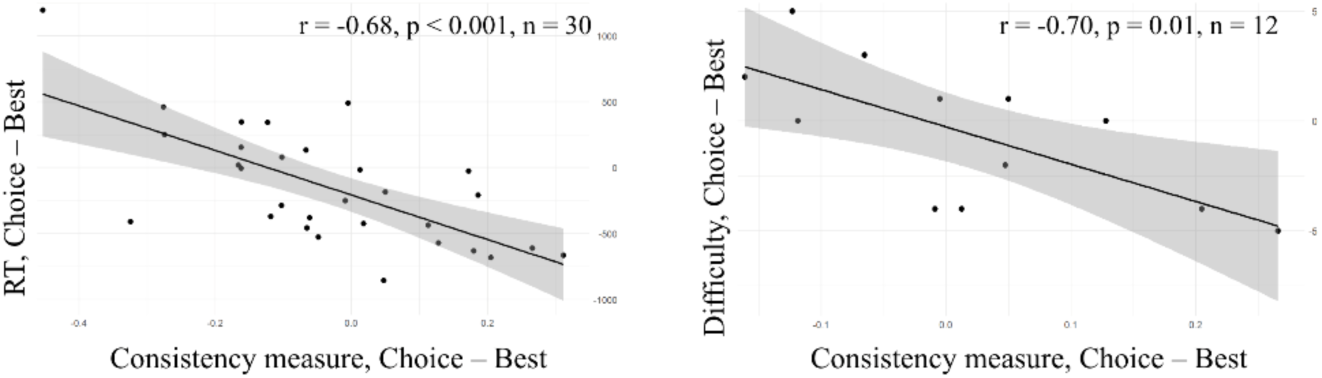
Correlation of consistency measure (Choice – Best) with self-reported difficulty ratings (Choice – Best) and RT of the decisions during the tasks (Choice – Best).

In terms of agreement between the self-report and behavior, we found that the self-reported policy accounts for the behavior to a greater extent, has a higher posterior probability and a smaller (better) rank with respect to other policies in case of Best as compared to the Choice condition as shown in figure 5.

**Figure 5:**
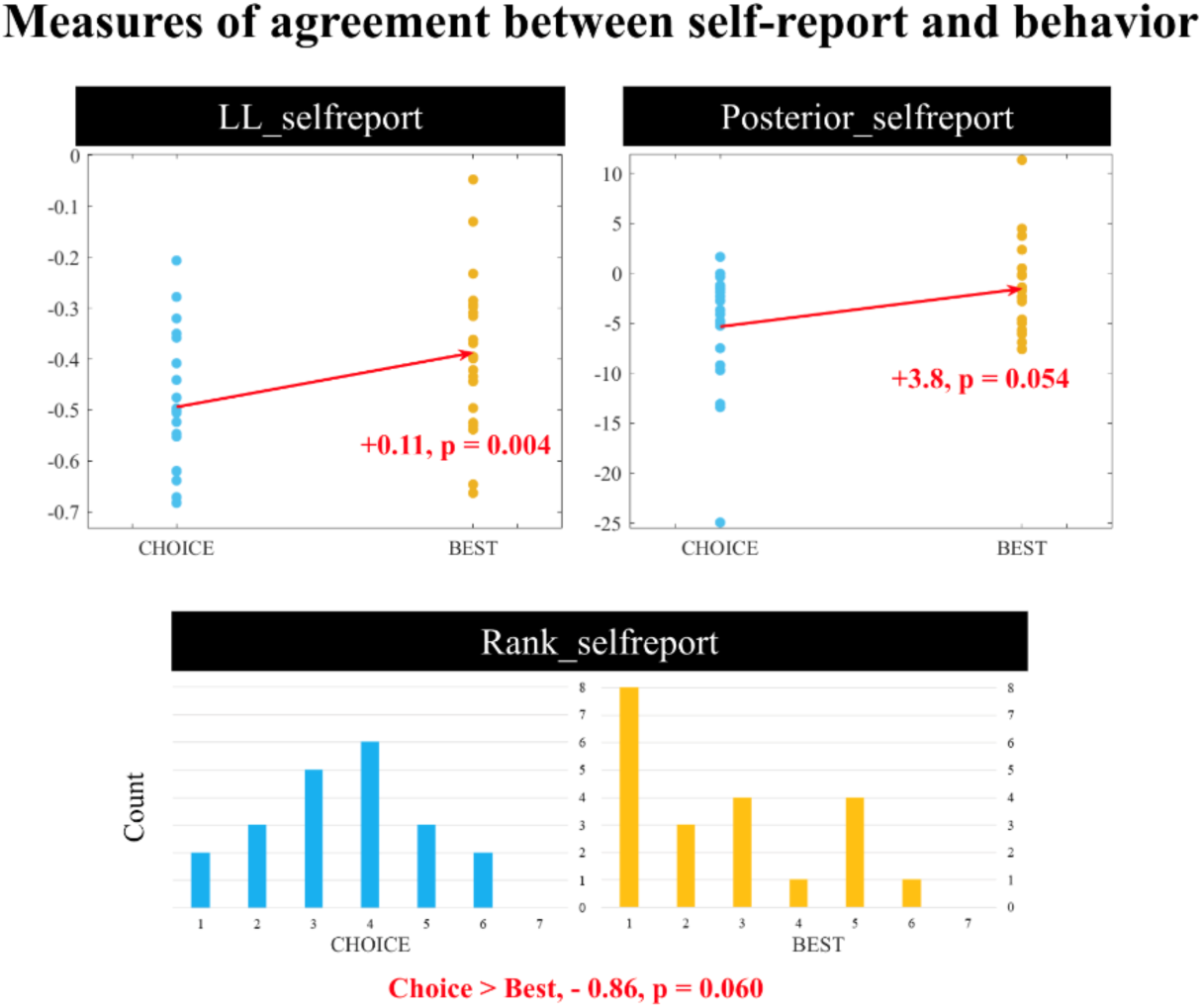
Measures of agreement between self-reported policies and behavior. The ‘self-reported policy’ here refers to the policy to which the responses to open-ended questions about task-strategy were mapped to. LL_selfreport – Comparison (paired, non-parametric) of the log-likelihood of the behavioral data given the self-reported policy; Posterior_selfreport – Comparison (paired, t-test) of the log-transformed posterior probability of the self-reported policy; Posterior_selfreport – Comparison (paired, non-parametric) of the rank of the posterior probability of the self-reported policy with respect to other candidate policies

There was a significant correlation across participants of the agreement measures with age and years of education. Measure LL_selfreport showed significant association with Age in Choice, such that there was an increase in agreement with increase in Age in the Choice condition. This was true for LL_selfreport in Choice relative to Best as well. (LL_selfreport and Age in Choice: Spearman rank correlation, r = 0.46, p = 0.032, n = 22; LL_selfreport, Choice – Best, and Age: r = 0.47, p = 0.03, n = 21).

Also, the relative agreement in Choice with respect to Best (Choice – Best) in terms of LL_selfreport (r = 0.38, p = 0.09, n = 21) has a positive trend with more years of education.

### fMRI results

A cluster in mid-cingulum (Figure 2E and Table 2) shows a positive effect of the main contrast, (Decision_Choice_ – Decision_Choice_control_) – (Decision_Best_ – Decision_Best_control_). Contrast values in this cluster are not correlated with differences in RT between the conditions across participants. A small volume correction of this contrast within an Anterior Cingulate Cortex (ACC) mask (WFU_PickAtlas 116; (Maldjian et al. 2003) showed a significant cluster in dorsal ACC (Figure 2F, Table 3) This region in dACC (figure 2F) shows a nearly significant correlation of the BOLD response at the time of decisions with the difference in consistency between the conditions. After controlling for RT, this relation becomes significant (Regression, *β_consistency_*_C−*B*_ = −1.34, *p* = 0.03, *n* = 30) We did not find any effects of the main BOLD contrast or of (Decision_Choice_ – Decision_Best_) in the ROI analysis of the middle orbito-frontal cortex (mOFC), but contrast values for (Decision_Choice_ – Decision_Best_) in this region show a positive association (r = 0.37, p = 0.046, n = 30) with the differences in consistency measure between the conditions (Figure 2G).

**Table 2:**
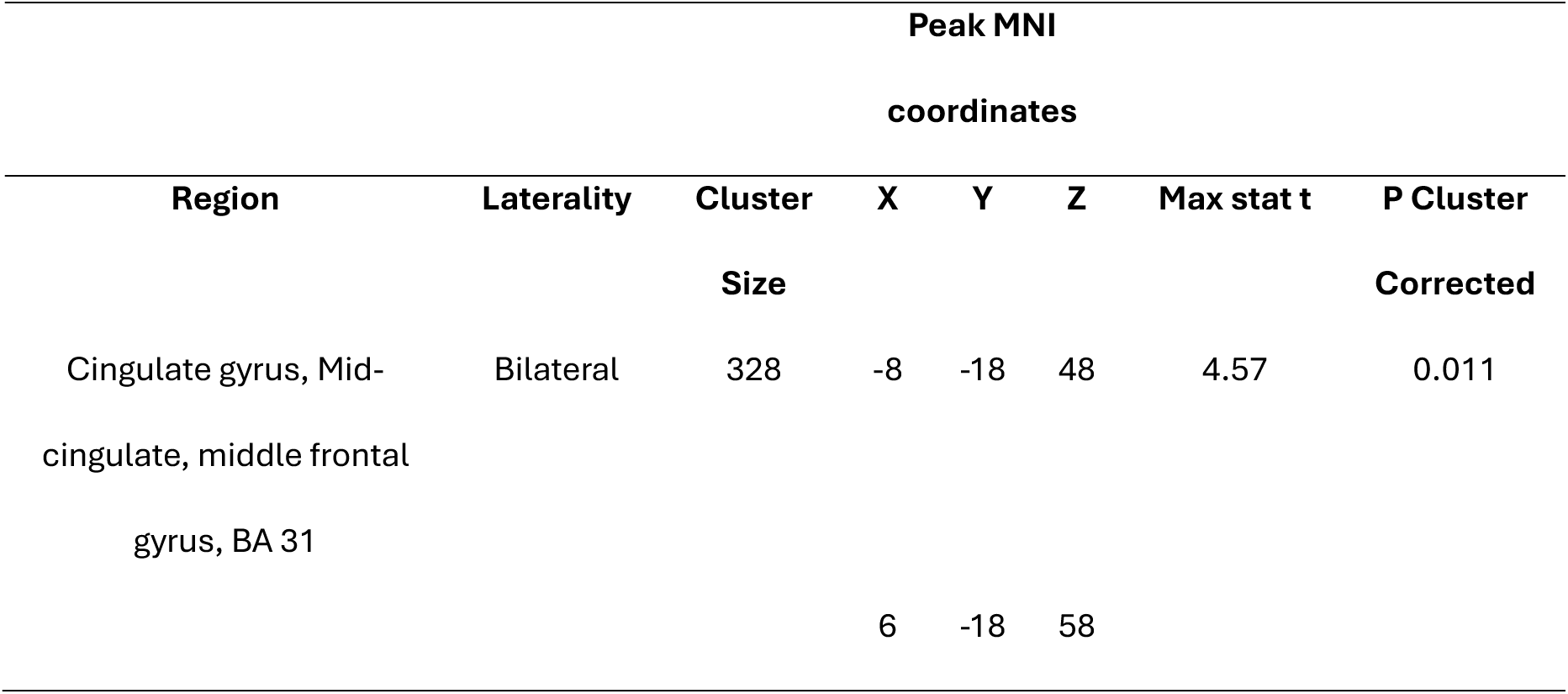
Cluster showing positive effect of the main contrast on BOLD response at the time of decision.

**Table 3:** Cluster showing significant effect of the task contrasts on BOLD response at the time of decision after small volume correction with anterior cingulate mask.

| Peak MNI |  |  |  |  |  |  |  |
| --- | --- | --- | --- | --- | --- | --- | --- |
| coordinates |  |  |  |  |  |  |  |
| Region | Laterality | Cluster | X | Y | Z | Max | P Cluster |
|  |  | Size |  |  |  | stat t | Corrected |
| **1. Small volume correction (ACC): main contrast** |  |  |  |  |  |  |  |
| Dorsal ACC | Bilateral | 41 | 4 | 18 | 28 | 4.05 | 0.04 |
|  |  |  | -4 | 14 | 28 |  |  |
| **2. Small volume correction (ACC): (Decision<sub>Choice</sub> – Decision<sub>Best</sub>)** |  |  |  |  |  |  |  |
| Dorsal ACC | Bilateral | 69 | 2 | 18 | 26 | 4.25 | 0.02 |
|  |  |  | -2 | 16 | 26 |  |  |

### Order effects

As described in the Methods section, there were nearly half (n = 16) of the participants who completed the Choice blocks first (Order 1; Group 1) and then the Best blocks while other half (n = 14) who completed the Best blocks first and then the Choice blocks (Order 2; Group 2). We assessed whether there was an effect of administering the tasks in a particular order on objectivity measures and fMRI results.

As shown in figure 2H, the consistency measure shows order effects, such for a task block that was second in the sequence, the consistency was 0.11 units greater on average than the first task (non-parametric paired samples test, p < 0.001). There were no order effects in the case of the non-randomness and distortion measures. These findings suggest that practice may have improved consistency.

In the case of BOLD responses, a two-sample t-test of the main contrast values in the significant cluster in mid-cingulum (Cluster in Figure 2E or Table 2) revealed that there was no significant difference between the groups (p = 0.814) indicating that the main effect was not driven by just one of the groups. Similarly, for the (Decision_Choice_ – Decision_Best_) contrast in the mid-cingulate region (Figure 2E), the above test did not reveal any significant differences (p = 0.508).

Correlation between the main contrast values in the mid-cingulate region above and the objectivity measure of consistency, however, showed order effects such that the association between the contrast values and the difference in consistency measure between Choice and Best was stronger and significant for group 2 (Figure 2E). We performed multiple regression of the contrast values in group 2, with differences in consistency and in RT between the conditions and the effect of consistency differences on contrast value remain significant in this analysis as well (Regression, *β_consistency_*_C−*B*_ = −0.90, *p* = 0.03, *n* = 14). In terms of alignment with specific policies, behaviorally policy 2 was employed to a greater extent in Best than Choice and hence, we also looked at whether behavioral alignment with policy 2 explained any variance in contrast values across the participants. We did not see any significant relation between the contrast values for group 2 and alignment with policy 2 after controlling for RT. Here behavioral alignment with policy 2 was quantified as log-likelihood of the behavioral data for policy 2. Note that this is same as the ‘Optim_P’ measure described earlier. Similarly, none of the other objectivity measures showed a significant linear relation with contrast values.

We also conducted a whole-brain exploratory two-sample t-test for the main and the (Decision_Choice_ – Decision_Best_) contrast, which revealed a cluster in posterior cingulate showing greater contrast values order 2 > order 1 in case of both the contrasts (Table S7, Figure S5).

## Discussion

In this study, we were interested in the neural and behavioral underpinnings of motivational subjectivity. In a within-subject design, we compared motivationally objective vs subjective decision-making by examining the behavior, self-report, and fMRI BOLD data from healthy participants performing a risky decision-making task designed to elicit both types of decision-making. The ‘Best’ condition was aimed at motivating more objective decision-making - as the choices were rewarded only when they were objectively the best on a per-trial basis, in terms of the points associated with the options; the task penalized choices that turned out worse than the unchosen option. This way, the reward structure aimed at incentivizing decisions based on externally provided or the objective goal of per-trial point maximization. The ‘Choice’ condition was more unconstrained in its reward structure wherein participants received the points associated with their choice irrespective of how it compared to the unchosen option, wherein one was freer to exercise subjective preferences.

The task worked as expected, revealing differences in policy/strategy between the Best and Choice conditions. Behavioral results showed that participants employed the objective policy to a greater extent in the Best task than the Choice task. For Best, the objectively optimal policy, corresponding to externally enforced goal of per-trial reward maximization, was policy 2 (Table S3) and this accounted for the participants’ behavior to the greatest extent (Figure 2A). For the Choice condition, objectively optimal policies were policies 1 or 2, depending on whether one assumes the goal to be long-term or per-trial reward maximization, respectively. As shown in figure 2A, participants’ data in Choice task is best explained by policy 5, which is neither of the above. At the level of individual participants too, objectively optimal policies accounted for the behavioral data to a significantly greater extent in the Best than the Choice condition (‘Optimal_L’ and ‘Optimal_P’, in figures 3B). While this supports that participants were more objective in Best than Choice in terms of applying objectively optimal policies, this approach assumes that the theoretically/objectively optimal policies above were conceptually accessible to all participants.

However, as can be seen in figure 3, 8 (out of 29) participants rated their familiarity with the concept of EV (pertinent to policy 1) as ‘1’, that is, ‘completely unaware of the concept’ and a little more than half the participants rated it as 5 or lower. This shows that relying on behavioral display of specific objective policies may not be an accurate measure of motivationally objective decisions. The rationale being that even if one cannot find the objectively optimal solution (or aren’t aware of it), they can still be objective within the set of solutions they do find or know.

Therefore, we also took a novel perspective on quantifying objectivity to assess task-manipulation. We expected there to be a greater consensus, across the participants, in employed policies in the Best (Objective) condition as it demanded following an externally-specified goal and relying solely on information available to all participants. We expected a greater variety in the policies employed in the Choice (Subjective) condition as it was less constrained externally. Our observations matched this expectation, both behaviorally and in self-reports. Behaviorally, across the participants, the posterior probability was largely localized to a single policy in the ‘Best’ condition, while it was more distributed in the ‘Choice’ condition, with at least two policies sharing the posterior probability mass (Figure 2A). Self-reported task strategies also mapped onto a greater variety of candidate policies in the Choice (Figure 2C). Additionally, from the self-reports, participants reported relying on subjective considerations like “intuition/gut-feeling” to a greater extent in the Choice condition. In the ratings for the importance of gamble characteristics, there was also more consistency across the subjects in how important the specific gamble characteristics were in the Best condition (Figure 2C), supporting that there was a greater consensus across participants in the applied policy in the Best condition. To assess whether a similar behavioral pattern existed at the level of individual participants, we defined a consistency measure to index commitment to a single policy or less noisy behavior. This measure reflected the consistency with which the winner policy, from the model-comparison at the level of individual participants, was applied throughout the task – irrespective of what policy it was. The rationale was that since the external or objective information was not changed across the trials of a task, if an individual switches between different policies or shows noisy behavior, leading to smaller consistency, it would indicate higher subjective influences. Consistency was not significantly different between the conditions (‘Consistency’ in figure 2B), however, task-induced differences in consistency may have been washed out by the task order effect, where the consistency sharply increased for second task irrespective of what task it was. Note that the participants performed two blocks of the same task first, followed by two blocks of the other later (order 1: Choice (x 2) ➔ Best (x 2); order 2: reverse of order 1). Please see supplementary discussion for more on task order effects and objectivity measures. But the effect of the task conditions on consistency was isolated by performing a between-subjects comparison of the consistency measure during the Best block for subjects for whom Best task was conducted first with the consistency during the Choice block for subjects for whom the Choice task was conducted first. This showed that the consistency in Best was significantly higher than the consistency in Choice (Figure 2H), indicating that, even at the level of individual participants, objective decision-making condition induced a greater commitment to a single policy, suggesting more objectivity.

Further, we found a greater agreement between self-report and behavior in objective (Best) than the subjective (Choice) decision-making condition in terms of the extent to which self-reported policy accounted for the behavior indicated by our measure ‘LL_selfreport’. This can also be seen by comparing figures 2A and 2C, where behavioral inference from the data pooled across the participants and the distribution of self-reported strategies are in agreement for the Best condition, with policy 2 having the highest posterior probability as well as the highest count. This wasn’t the case in the Choice condition however, where the posterior probability was highest for policy 5, followed by policy 2, but in self-report, it was unequivocally policy 2 that had the highest count. A higher agreement between the self-report and behavior in the objective condition may arise due to several reasons, such as: (1) Higher self-awareness: The behavior in the objective condition was influenced only by the processes that were consciously accessible (hence, could be reported). Ericsson and Simon (1980) argued that inaccuracy in self-reports often results from not directly heeding to the queried information during the task. (2) Higher execution accuracy: Execution of the intended (self-reported) policy was accurate and without mistakes in the objective condition, resulting in the intended policy accounting for the behavioral data to a greater extent. (3) Higher self-report accuracy: Higher proficiency or willingness to accurately recall or verbalize the policies employed in the objective condition. Brenner and DeLamater 2016; Bernstein, Chadha, and Montjoy (2001) show that in case of normative behaviors like voting, exercising, etc, individuals tend to inaccurately self-report either to appear a certain way to interviewers or to themselves. However, the simplicity of the self-reported policy (such as, policy 6 being mathematically simpler to execute than policy 1) and the agreement measures did not show a significant association. Further, since it was a within-subject study, if an individual was motivated to provide inaccurate or normative reports of the task-strategy, it is reasonable to assume that it must be so in both conditions. While we cannot rule out these possibilities, based on the above, the case for there being higher self-awareness in the Best/objective decision-making seems stronger than the one for this condition having higher execution accuracy or higher self-report accuracy. We also found that with increasing age or increase in years of education, there was an increase in the agreement measure in subjective (Choice) decision-making condition, relative to the objective condition. Absolute agreement in Choice also increased with age. This may be driven by the age or education leading to greater awareness of one’s subjective decision-making. Alternatively, these factors could also lead to greater willingness to accurately report one’s decision strategies or better execution accuracy.

We also found that the objective (Best) condition was characterized by a greater RT. Since the objective condition was associated with greater consistency, which we may reflect narrower consideration of alternative policies, and consistency was negatively associated with RT, one would expect the objective condition to have lower RT. However, a greater decision RT in the objective condition could result from reasons beyond consideration of alternative policies, like the time taken in implementing the policy, that is, evaluating the gamble and the sure thing as per the policy.

There was a greater BOLD response in the mid-cingulum (Figure 2E) at the time of decision in Choice (relative to Control) than in Best (relative to Control) condition, suggesting its active involvement to a greater extent in subjective decision-making. Since there were significant differences in RT between the conditions, we looked at the relation between the values of the main contrast above in this region and RT differences, across subjects. This yielded no significant relation between the two, suggesting against the possibility that the contrast effect in this region cannot simply be accounted by time on task (Grinband et al. 2011; Brown 2011). The effect, therefore, is likely to be driven by the cognitive differences between objective and subjective decision-making.

As discussed earlier, there was a smaller consistency and a greater variety of policies employed in the subjective condition and we wondered whether these were associated with the BOLD contrast effects. Across participants, we found a negative but non-significant relation between the main contrast values in this region and differences in consistency between the conditions. This relation was significantly negative for the participants (group 2) who performed the Choice task after the Best task, such that the BOLD signal in mid-cingulum was relatively smaller in subjective decision making when there was a larger relative consistency in this condition. It is currently unclear why the relation between contrast values and consistency would show order effects, but these may be driven by an interaction effect of fatigue or familiarity and the task condition on consistency. This suggests that this region may be involved in processes relating to generating alternatives for policies or choosing between them. This does not seem to be involved in selecting specific choice options as the differences in choice probabilities did not show any significant relation or trend with contrast values. As noted earlier, another difference between the conditions was that the behavior in subjective condition was accounted by multiple policies, while in objective, it was accounted largely by policy 2. However, the differences in alignment with policy 2 between the conditions did not account for the contrast value suggesting that the BOLD effect was not driven by differences in extent to which policy 2 was employed.

The relation with consistency measure is aligned with the possibility that the cluster in mid-cingulum/PCC may be involved in generating options for policies or choosing between them. A greater BOLD signal in this region may reflect a broader consideration of alternative policies or a greater conflict between them, manifesting as smaller consistency. Previous work has implicated regions in mid- and posterior-cingulate in subjective valuations (Smith, Ahern, and Lane 2019; Kable and Glimcher 2007; Sripada et al. 2011; Fitzgerald et al. 2010; Greene et al. 2001). Our result shows that these regions may also be involved in subjectively choosing between different policies or evaluation criteria. This interpretation is broadly in line with prior work implicating cingulate regions in conflict monitoring and control allocation (Botvinick et al., 2001; Shenhav, Botvinick, & Cohen, 2013), and posterior cingulate cortex in adaptive behavioral shifts and evaluating alternative courses of action in changing environments (Pearson et al. 2011; Leech and Sharp 2014)

Further, small volume correction of the (Decision_Choice_ – Decision_Best_) contrast with anterior cingulate cortex (ACC), revealed significant effects in dorsal ACC (dACC), such that there was a greater BOLD signal in this region at the time of decision in subjective decision-making condition. This region shows a weaker (still significant) effect of the main contrast. Further, (Decision_Choice_ – Decision_Best_) contrast values in dACC were negatively associated with differences in consistency between the conditions, even after controlling for RT. Similar to mid-cingulate and PCC, this also supports the interpretation that dACC may be involved in generating alternatives for or deliberating over candidate strategies. This is in-line with previous work (Walton, Devlin, and Rushworth 2004; Guggisberg et al. 2007), especially with Shenhav and Buckner (2014) showing greater BOLD signal in a region in dACC during choice was associated with a greater likelihood choice reversal later on.

We also performed small volume correction of exploratory fMRI analysis with an orbito-frontal gyrus mask as well as performed an ROI analysis of the contrasts with bilateral middle orbitofrontal cortex (mOFC) and neither found the effect of main contrast nor (Decision_Choice_ – Decision_Best_) contrast in this region. However, this showed a significant association with the consistency measure across the participants such that when the consistency was greater, the BOLD signal at the time of decision was greater. This may suggest the active involvement of this region in maintaining a policy or preventing generation of candidate policies or switching to them. Interestingly, it is not associated with the measure of alignment with optimal policies. This provides an interpretation of its function that may tie together the results from previous studies, with some implicating it in rational behaviors (Bechara et al. 1994; De Martino et al. 2006) while some in subjective valuations/decisions (Koenigs and Tranel 2008; Grabenhorst and Rolls 2009). This work suggests that it may not necessarily be associated with specific policies, objective or subjective, but with maintaining a given goal/policy. This interpretation is supported by Fellows (2011) discussing various lesion studies that suggest that damage to OFC or vmPFC is associated with inconsistent preference judgments, and by Zarr and Brown (2023) implicating it in goal representation.

## Conclusion

In this study, we investigated the neural and behavioral basis of motivationally objective versus subjective decision-making using a within-subject risky decision-making task. The results suggest that the task manipulation was effective in eliciting more objective decision-making in the Best condition and more subjective decision-making in the Choice condition. Behaviorally, decision-making in the motivationally subjective condition showed smaller alignment with objectively optimal policies, smaller consensus across participants in the applied policies, smaller consistency in the employed policy by individual participants, smaller agreement between self-report and behavior, and smaller RT relative to the Choice condition. We also present an approach to studying motivational subjectivity that does not rely on deviations from a single theoretically optimal policy such as EV maximization and highlight a drawback of the latter approach in that participants varied considerably in their familiarity with the concept of expected value, which may confound the extent of displayed subjectivity, as measured by deviation from the EV policy. At the neural level, motivationally subjective, relative to objective, decision-making was associated with greater BOLD response in mid-cingulum/posterior cingulate cortex and dorsal anterior cingulate cortex at the time of decision, with these effects relating negatively to behavioral consistency, suggesting involvement in generating alternatives for decision policies or choosing between them. In contrast, activity in middle orbitofrontal cortex was associated with greater consistency without showing any significant differences between the conditions, suggesting a possible role in maintaining an adopted policy rather than representing a specific objective or subjective strategy.

## Limitations

It is important to note there was one more difference between the task conditions, other than the task-manipulation, that could have potentially led to the main effect. There was more predictability of the decision outcome in-built in Choice than the Best condition. This is because if say the sure thing was chosen in Choice, then the participant had certainty about what the outcome would be; it would be equal to the value associated with sure thing. But in the case of Best, upon choosing any option, including the sure thing option, there is uncertainty about the outcome as the outcome of the trial depended on a comparison of the outcomes from both the options. But if this indeed has an effect, then one must see an association of contrast values with how often the ST was chosen, however, there was nearly zero correlation between the main contrast values and the differences in choice probability for ST. This suggests that the effect of the main contrast in mid-cingulum above may not be (linearly) associated with the greater certainty about the outcome during decisions in Choice condition with respect to the Best. The correlation of the main effect with objectivity measures establishes differences in objectivity as one of the drivers of the main effect. So, the above may not have entirely driven the main effect but could have contributed to it. Further, in future studies, the goal of our study could be better achieved by including two or more conditions for the objective decision-making task that reinforce different task strategies. Together, these conditions would provide a better control for subjective decision-making, allowing the neural correlates of subjectivity to be isolated beyond the implementation of specific policies.

## Acknowledgments

We would like to thank the staff of the imaging research facilities at Indiana University for their assistance with fMRI data collection.

## References

American Psychological Association. (n.d.). Objectivity. In APA dictionary of psychology. Retrieved July 1, 2026, from https://dictionary.apa.org/objectivity.

Bechara, A., Damasio, A. R., Damasio, H., & Anderson, S. W. (1994). Insensitivity to future consequences following damage to human prefrontal cortex. Cognition, 50(1–3), 7–15.

Bechara, A., Damasio, H., Tranel, D., & Damasio, A. R. (1997). Deciding advantageously before knowing the advantageous strategy. Science, 275(5304), 1293–1295.

Bernstein, R., Chadha, A., & Montjoy, R. (2001). Overreporting voting. Public Opinion Quarterly, 65(1), 22–44.

Brandstätter, E., Gigerenzer, G., & Hertwig, R. (2006). The priority heuristic: Making choices without trade-offs. Psychological Review, 113(2), 409–432.

Brenner, P. S., & DeLamater, J. (2016). Lies, damned lies, and survey self-reports? Identity as a cause of measurement bias. Social Psychology Quarterly, 79(4), 333–354.

Brown, J. W. (2011). Medial prefrontal cortex activity correlates with time-on-task: What does this tell us about theories of cognitive control? NeuroImage, 57(2), 314–315.

De Martino, B., Kumaran, D., Seymour, B., & Dolan, R. J. (2006). Frames, biases, and rational decision-making in the human brain. Science, 313(5787), 684–687.

Ericsson, K. A., & Simon, H. A. (1980). Verbal reports as data. Psychological Review, 87(3), 215– 251.

Fellows, L. K. (2011). Orbitofrontal contributions to value-based decision making: Evidence from humans with frontal lobe damage: The human frontal lobes in choice and learning. Annals of the New York Academy of Sciences, 1239(1), 51–58.

Fitzgerald, T. H. B., Seymour, B., Bach, D. R., & Dolan, R. J. (2010). Differentiable neural substrates for learned and described value and risk. Current Biology, 20(20), 1823–1829.

Grabenhorst, F., & Rolls, E. T. (2009). Different representations of relative and absolute subjective value in the human brain. NeuroImage, 48(1), 258–268.

Greene, J. D., Sommerville, R. B., Nystrom, L. E., Darley, J. M., & Cohen, J. D. (2001). An fMRI investigation of emotional engagement in moral judgment. Science, 293(5537), 2105–2108.

Grinband, J., Savitskaya, J., Wager, T. D., Teichert, T., Ferrera, V. P., & Hirsch, J. (2011). The dorsal medial frontal cortex is sensitive to time on task, not response conflict or error likelihood. NeuroImage, 57(2), 303–311.

Guggisberg, A. G., Dalal, S. S., Findlay, A. M., & Nagarajan, S. S. (2007). High-frequency oscillations in distributed neural networks reveal the dynamics of human decision making. Frontiers in Human Neuroscience, 1, 14.

Hsu, M., Bhatt, M., Adolphs, R., Tranel, D., & Camerer, C. F. (2005). Neural systems responding to degrees of uncertainty in human decision-making. Science, 310(5754), 1680–1683.

Hsu, M., Krajbich, I., Zhao, C., & Camerer, C. F. (2009). Neural response to reward anticipation under risk is nonlinear in probabilities. The Journal of Neuroscience, 29(7), 2231–2237.

Hutcherson, C. A., Montaser-Kouhsari, L., Woodward, J., & Rangel, A. (2015). Emotional and utilitarian appraisals of moral dilemmas are encoded in separate areas and integrated in ventromedial prefrontal cortex. The Journal of Neuroscience, 35(36), 12593–12605.

Jeffreys, H. (1961). Theory of probability (3rd ed.). Oxford University Press.

Kable, J. W., & Glimcher, P. W. (2007). The neural correlates of subjective value during intertemporal choice. Nature Neuroscience, 10(12), 1625–1633.

Kahneman, D., & Tversky, A. (1979). Prospect theory: An analysis of decision under risk. Econometrica, 47(2), 263.

Kass, R. E., & Raftery, A. E. (1995). Bayes factors. Journal of the American Statistical Association, 90(430), 773.

Koenigs, M., & Tranel, D. (2008). Prefrontal cortex damage abolishes brand-cued changes in cola preference. Social Cognitive and Affective Neuroscience, 3(1), 1–6.

Leech, R., & Sharp, D. J. (2014). The role of the posterior cingulate cortex in cognition and disease. Brain, 137(Pt 1), 12–32.

Maldjian, J. A., Laurienti, P. J., Kraft, R. A., & Burdette, J. H. (2003). An automated method for neuroanatomic and cytoarchitectonic atlas-based interrogation of fMRI data sets. NeuroImage, 19(3), 1233–1239.

Modak, P., & Brown, J. W. (2026). Neural correlates of the subjective experience of free-will during value-based risky decisions: A pilot study. bioRxiv. 10.64898/2026.07.12.738094.

Modak, P., Hutslar, C., Polk, R., Atkinson, E., Fisher, L., Macy, J., Chassin, L., Presson, C., Finn, P. R., & Brown, J. W. (2021). Neural bases of risky decisions involving nicotine vapor versus monetary reward. NeuroImage: Clinical, 32, 102869.

Park, S. Q., Kahnt, T., Rieskamp, J., & Heekeren, H. R. (2011). Neurobiology of value integration: When value impacts valuation. The Journal of Neuroscience, 31(25), 9307–9314.

Paulus, M. P., & Frank, L. R. (2006). Anterior cingulate activity modulates nonlinear decision weight function of uncertain prospects. NeuroImage, 30(2), 668–677.

Payne, J. W., Bettman, J. R., Coupey, E., & Johnson, E. J. (1992). A constructive process view of decision making: Multiple strategies in judgment and choice. Acta Psychologica, 80(1–3), 107– 141.

Pearson, J. M., Heilbronner, S. R., Barack, D. L., Hayden, B. Y., & Platt, M. L. (2011). Posterior cingulate cortex: Adapting behavior to a changing world. Trends in Cognitive Sciences, 15(4), 143–151.

Rao, L.-L., Li, S., Jiang, T., & Zhou, Y. (2012). Is payoff necessarily weighted by probability when making a risky choice? Evidence from functional connectivity analysis. PLOS ONE, 7(7), e41048.

Rao, L.-L., Zhou, Y., Xu, L., Liang, Z.-Y., Jiang, T., & Li, S. (2011). Are risky choices actually guided by a compensatory process? New insights from fMRI. PLOS ONE, 6(3), e14756.

Shenhav, A., Botvinick, M. M., & Cohen, J. D. (2013). The expected value of control: An integrative theory of anterior cingulate cortex function. Neuron, 79(2), 217–240.

Shenhav, A., & Buckner, R. L. (2014). Neural correlates of dueling affective reactions to win-win choices. Proceedings of the National Academy of Sciences of the United States of America, 111(30), 10978–10983.

Smith, R., Ahern, G. L., & Lane, R. D. (2019). The role of anterior and midcingulate cortex in emotional awareness: A domain-general processing perspective. Handbook of Clinical Neurology, 166, 89–101.

Sripada, C. S., Gonzalez, R., Phan, K. L., & Liberzon, I. (2011). The neural correlates of intertemporal decision-making: Contributions of subjective value, stimulus type, and trait impulsivity: Neural correlates of intertemporal decision-making. Human Brain Mapping, 32(10), 1637–1648.

Valentin, V. V., Dickinson, A., & O’Doherty, J. P. (2007). Determining the neural substrates of goal-directed learning in the human brain. The Journal of Neuroscience, 27(15), 4019–4026.

Venkatraman, V., Payne, J. W., Bettman, J. R., Luce, M. F., & Huettel, S. A. (2009). Separate neural mechanisms underlie choices and strategic preferences in risky decision making. Neuron, 62(4), 593–602.

Walton, M. E., Devlin, J. T., & Rushworth, M. F. S. (2004). Interactions between decision making and performance monitoring within prefrontal cortex. Nature Neuroscience, 7(11), 1259–1265.

Wang, G., Li, J., Wang, P., Zhu, C., Pan, J., & Li, S. (2019). Neural dynamics of processing probability weight and monetary magnitude in the evaluation of a risky reward. Frontiers in Psychology, 10, 554.

Zarr, N., & Brown, J. W. (2023). Foundations of human spatial problem solving. Scientific Reports, 13(1), 1485.

