## Supplementary Material for "Motivationally objective versus subjective decision-making: Neural correlates, behavior, and self-report"

### I. Supplementary Methods

#### 1. Demographics

n = 30

*Table S1: Demographic details of the participants*

| Sex | Race | Age (years) | Education (years) |
|---|---|---|---|
| Female = 18<br>Male = 12 | Caucasian = 17<br>Asian Descent = 9<br>Other = 4<br>Black or African American = 0<br>American Indian/Alaska Native = 0<br>Native Hawaiian/Pacific Islander = 0 | Mean = 22<br>Standard deviation = 4.7<br>Maximum = 42<br>Minimum = 18 | Mean = 14.6<br>Standard deviation = 2.4<br>Maximum = 21<br>Minimum = 10 |

Age was not significantly correlated with years of education (r = 0.18, p = 0.346).

#### 2. Gambles

*Table S2: Details of the gamble options*

| Gambles | Values (V1 > V2 > V3) | | Probabilities | |
|---|---|---|---|---|
| **1** | V1 | 50 | P1 | 0.2 |
| | V2 | 15 | P2 | 0.5 |
| | V3 | -20 | P3 | 0.3 |
| **2** | V1 | 64 | P1 | 0.2 |
| | V2 | 10 | P2 | 0.6 |
| | V3 | -10 | P3 | 0.2 |
| **3** | V1 | 38 | P1 | 0.5 |
| | V2 | -7 | P2 | 0.5 |
| | V3 | NA | P3 | NA |
| **4** | V1 | 40 | P1 | 0.5 |
| | V2 | 30 | P2 | 0.2 |
| | V3 | -10 | P3 | 0.3 |
| **5** | V1 | 38 | P1 | 0.3 |
| | V2 | 8 | P2 | 0.45 |
| | V3 | -1 | P3 | 0.25 |

#### 3. Certainty equivalent task

We developed a webpage using JavaScript, HTML, CSS, and PHP, hosted on our lab's server, to conduct the certainty equivalent task (Paulus and Frank 2006). A personalized link was sent to the participants, where for each gamble, they chose between the gamble and a sure-thing for a range of sure thing values. Their choice responses were fitted using a logistic regression model for each of the five gambles, which was used to find sure-thing values corresponding to choice probabilities, 0.35 and 0.65, referred to as $ST_{35}$ and $ST_{65}$.During a task block, for any trial, we first randomly selected the gamble and then randomly selected one of the two sure-thing values corresponding to it with an added Gaussian noise ($\mu = 0$). The standard deviation was dependent on how steep the slope of the logistic model was around the $ST_{35}$ and $ST_{65}$ values such that it was equal to the average deviation in ST values required for a deviation of decision probability of 0.2 around $ST_{35}$ and $ST_{65}$. So, for a steeper slope, the standard deviation of the Gaussian used for adding the noise was smaller.

#### 4. Payment

Participants received $10 for completing the certainty equivalent task (performed online, remotely) and $30/hour for the fMRI session. They also received a $10 bonus for arriving on time for the experiment.

Participants were informed that they could also earn from the tasks, proportional to the points they earned during each of them:

- 0 to $5 for the certainty equivalent task
- 0 to 5$ for the choice task
- 0 to 5$ for the best task
- 0 to 2$ for the control task

Payment was made in cash.

For the certainty equivalent and choice task, their total points as a proportion of 1000 points was used to find the proportion of $5 they were paid in each task; if they earned more than or equal to 1000, they were paid the full $5 in each task. In these tasks, there was no concept of being 'correct' and hence, we arbitrarily chose 1000 points as our reference for converting points to money. In the control task, the proportion of $2 that they were paid was equal to the proportion of correct trials. In best task, participants were paid a proportion of $5 that was equal to the proportion of 'm' trials that were correct, where m was 70% of the total number of trials. This was done because it was hard to earn points in the Best task and we wanted the payment to be less stringent than the control.

#### 5. Predicting scanner start-time

**The problem: Find x**

It is essential to know, for analysis, the time at which scanner began collecting the fMRI BOLD data with respect to the start of the behavioral task in an event-based design. However, for a subset of subjects (n = 22, group U), we did not have this scanner start time due to a technical issue in recording it. We had this information for some subjects (n = 8, group K) however. We refer to these as groups U and K, where group U are the subjects whose scanner start time was unknown and for group K it was known.

Essentially, the problem was to determine ‘x’ in secs for every run of a particular subject which is shown in figure S1. From the task, we knew that ‘x’ could be any of {2,4,6,8} for the task blocks (figure S1A) and {5.14, 7.14, 9.14, 11.14, 13.14, 15.14, 17.14} for the control blocks (figure S1B) and for a run, we used a trained SVM to select between these possible values of x.

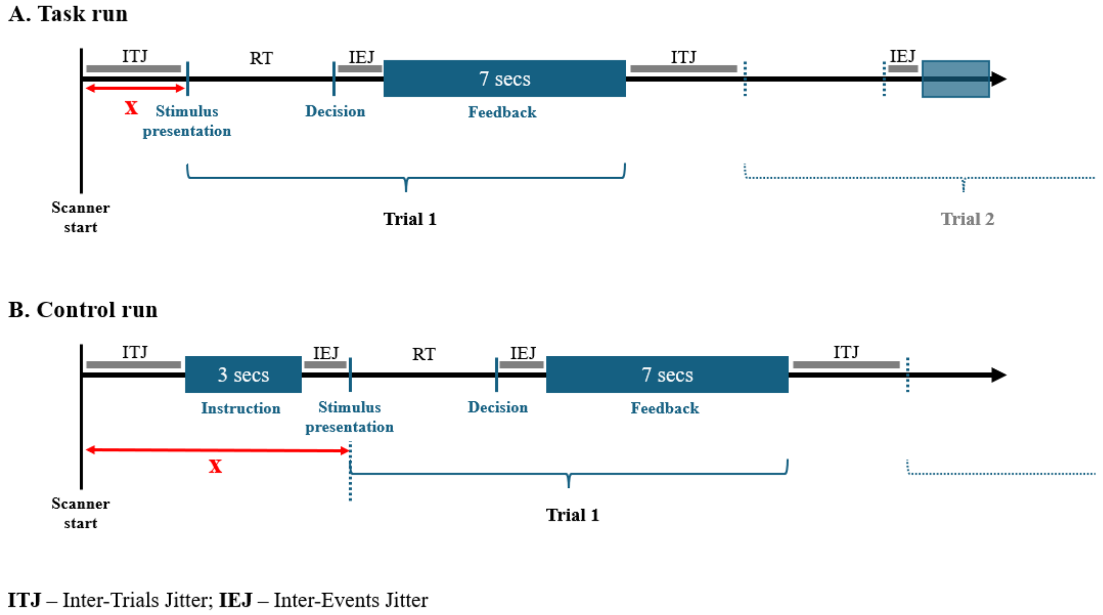

*Figure S1: The unknown interval, X, in task (A) and control (B) runs for group U. We knew x for some participants whom we refer to as group K. For some participants, we did not know the value of x and refer to them as group U. We used a machine learning approach to predict x for group U by training a support vector machine on data from group K. ‘x’ could be any of {2, 4, 6, 8} secs in A and any of {5.14, 7.14, 9.14, 11.14, 13.14, 15.14, 17.14} secs in B.*

**Training data:** For each participant in group K, we found the whole brain beta maps for the ‘Dec’ and ‘Not Dec’ event regressors using general linear modeling in SPM12. For the task conditions (Both, Choice and Best), there was one ‘Dec’ regressor representing the time when a decision was submitted and seven ‘Not Dec’ regressors representing various timepoints during a trial which were not the timepoints when a decision was submitted. Similarly, for the control blocks, there was one ‘Dec’ regressor and 13 ‘Not Dec’ regressors. This was the training data. There are several other ways to generate training data as described in the following paragraph, but the aforementioned way turned out the best in terms of cross-validation accuracy.

**Hyper-parameters, training, and LOSO-CV:** SVM with best hyper parameters trained to classify a beta map in left pre and post-central gyrus as 'Dec' and 'Not Dec'.

The beta-maps for 'Dec' and 'Not Dec' were masked to isolate the left precentral and postcentral gyrus. We used AFNI's 3dsvm to train a support vector machine (SVM) to classify these masked beta maps as 'Dec' and 'Not Dec' using the training data from group K (8 subjects). To test the accuracy of the classifier, we performed a leave-one-subject-out cross-validation (LOSO-CV). We trained the SVM on data from 7 subjects while using the data from the remaining subject for testing and repeated this 8 times so that a different subject is used for testing every time. The 'testing' was performed in the following way: For a particular run of the test subject, a GLM with same regressors as stated in the section 'Training data' was estimated for each possible delay (x). For a task run, four GLMs were estimated corresponding to each possible delay {2,4,6,8}. Similarly, for a control run, seven GLMs were estimated corresponding to its possible delay times. So, for a subject and a particular run, we had a 'Dec' beta map from left precentral and postcentral gyrus for each of the possible delay. we then used the trained SVM to classify these beta maps as 'Dec' or 'Not Dec'. Specifically, a score was generated for each of the beta maps indicating how far it was from the classification boundary and on which side. The delay time that was farthest from the boundary and on the side of 'Dec' was taken as the x for that particular run of that subject. We already knew the ground truth 'x' for the runs of the test subject and compared them with the predicted x values to evaluate the accuracy of the prediction.

We looked at this accuracy for different hyper-parameters like different ways of generating training data (different number of regressors or at different time points) or using the SVM of a different complexity (default or 100) and degree (1 or 3). The best cross-validation accuracy was 79%. This was attained with the SVM at its default complexity with a degree-3 kernel and the training data generated in a manner described above. This was also achieved with a kernel of degree 1 but we chose to go ahead with the kernel of degree 3 over that because in the latter case the inaccuracies were better spread across the subjects and the type of runs.

We now trained the SVM with the best hyper-parameters determined above on the training data from all 8 subjects which I will refer to as 'SVM_trained'.

**Finding x for group U:** For a particular subject in group U and for a particular run, SVM_trained was used to predict an x the same way as described in the previous section on how SVM was tested.

#### 6. Behavioral analysis and candidate behavioral policies

*Table S3: Candidate policies for performance in Choice and Best conditions. These are considered stochastic policies with 'Gamble selected' indicating more likely to select gamble, 'Indecisive' indicating random selection, and 'ST selected' indicating more likely to select sure thing. EVG – Expected Value of Gamble (P1V1 + P2V2 + P3V3). Please refer to figure 1B for notations of different gamble characteristics.*

| # | Policy[1] | Policy Name |
|---|---|---|
| 1 | $Gamble\ selected, if\ EVG > c * ST;$<br>$Indecisive, if\ EVG = c * ST;$<br>$ST\ selected, if\ EVG < c * ST$ | Expected Value |

| | | |
|---|---|---|
| 2 | $\boldsymbol{When\ V1\ >\ ST\ and\ V2 \geq\ ST},$<br>$Gamble\ selected, if\ P1\ +\ P2\ >\ P3;$<br>$Indecisive\ if\ P1 + P2 = P3;$<br>$ST\ selected, if\ P1\ +\ P2\ <\ P3$<br>$\boldsymbol{When\ V1\ >\ ST\ and\ V2 < ST},$<br>$Gamble\ selected, if\ P1\ >\ P2\ +\ P3;$<br>$Indecisive, is\ P1 = P2 + P3;$<br>$ST\ selected, if\ P1\ <\ P2\ +\ P3$<br>$\boldsymbol{When\ V1\ \leq\ ST},$<br>$Choose\ sure\ thing$ | Probability of delivering the better outcome |
| 3 | $Gamble\ selected, if\ P1 * V1 + P2 * V2 > c * ST;$<br>$Indecisive, if\ P1 * V1 + P2 * V2 = c * ST;$<br>$ST\ selected, if\ P1 * V1 + P2 * V2 < c * ST$ | Expected gains |
| 4 | $Gamble\ selected, if\ P1 * V1 > c * ST;$<br>$Indecisive, if\ P * V1 = c * ST;$<br>$ST\ selected, if\ P1 * V1 < c * ST$ | Expected value of the highest gamble outcome |
| 5 | $\boldsymbol{When\ V2 \geq\ ST},$<br>$Gamble\ selected, if\ P1 * V1\ +\ P2 * V2 > c * ST;$<br>$Indecisive, if\ P1 * V1 + P2 * V2 = c * ST$<br>$ST\ selected, if\ P1 * V1\ +\ P2 * V2 <\ c * ST$<br>$\boldsymbol{When\ V2 < ST\ and\ (V1 \geq ST\ or\ V1 < ST)},$<br>$Gamble\ selected, if\ P1 * V1 > c * ST;$<br>$Indecisive, if\ P1 * V1 = c * ST$<br>$ST\ selected, if\ P1 * V1 <\ c * ST$ | Expected value of the gamble outcomes higher than ST |
| 6 | $\boldsymbol{When\ V2\ >\ c * ST},$<br>$Gamble\ selected;$<br>$\boldsymbol{When\ V2 = c * ST};$<br>$Indecisive;$<br>$\boldsymbol{When\ V2 < c * ST\ and\ (V1 \geq c * ST\ or\ V1 < c * ST)},$<br>$ST\ selected$ | Number of gamble outcomes higher than ST |
| 7 | $Choose\ the\ option\ on\ the\ right$ | Random choice |

The gamble characteristics in table above are described in Figure 1B. Participants' choices were coded as '1' for sure thing and '0' for gamble. We performed logistic regression of the choice data with the regressor formed by subtracting the sure-thing value from the corresponding (combination of) gamble characteristic(s) for each of the policies in table S3 in both conditions. In the above, 'c' is a free parameter and accounts for subjective deviation from the candidate policies.

We did not include policies such as 'always choose ST' or 'always choose gamble' because even when the choice probability for either of the options was close to 1 for some gambles in some participants, their RT was not significantly different from the instances when the choice probability was close to 0.5. These RT values, when choices were on-sided, were also significantly more than the RT in the control blocks. This suggests the choices in these instances were not driven by such habitual policies.

We performed optimization of the parameters both, at the level of individual participants, and across the participants.

At the level of individual participant, the parameters for a policy were optimized using MATLAB's genetic algorithm to maximize the log-likelihood of the behavioral data from a particular participant and for a particular task condition. This way, we had optimal parameters (parameters c and α for policies 1 and 3-6, and parameter α for policy 2) for each policy, computed separately for each task condition per participant. For each participant and for a particular task condition, we performed model comparison using AIC between these policies at their optimal parameter values to identify a 'winner' policy.

For the analysis across the participants, we pooled together the behavioral data for all participants for a particular task condition and used Bayesian approach to identify the policy that was most likely given this data. Prior on the policies were such that it penalized the complexity in a way that was aligned with the idea of AIC.

This was done in the following way:

$$AIC_i = 2 * k_i - 2 * \ln(L_i) \tag{S1}$$

$Rearranging,$

$$AIC_i = 2(1 - \ln(L_i * e^{-k_i})) \tag{$S2$}$$

The idea of penalizing complexity in AIC is to multiply the likelihood of data given a policy/model with the factor $e^{-k_i}$ where $k_i$ represents number of free parameters in the policy. In the Bayesian approach, I incorporated this idea in the prior, such that the prior probability of the policy $i$ was taken as:

$$P(\pi_i) = \frac{e^{k_i}}{\sum_{i=1}^{m} e^{k_i}} \tag{S3}$$

$$P{:}\, probability, k_i{:}\, number\ of\ parameters, L_i{:}\, likelihood\ of\ the\ data,$$

$$i{:}\, policy\ index, m{:}\, numbe\ of\ policies$$

In calculating the marginal likelihood of the data for each policy, the prior over the parameter space was taken to be uniform as there was no reason to believe a particular parameterization to be more likely than the other. The parameter space was bounded with α ← [0 1] and c ← [0 3] and ten thousand points were sampled from it to find the marginal likelihood.

Additionally, policies 1 and policies 3-6 were individually optimized with parameters {α, c1} and policy 2 with parameter α to maximize the likelihood of the data pooled across the participants. In estimating posterior probabilities, what was used here was the likelihood of the data for each of the models at their optimal parameterization, instead of marginal likelihood (that is, the likelihood integrated across parameter space). Probability of data was taken as the weighted sum of these likelihoods, weighed by the prior probabilities. This is a measure of how much a policy at its optimum account for the data, relative to other policies. Please see the equations below for further clarity on the two analyses.

Analysis with integrated likelihood,

$$P(\pi_i \mid data) = \frac{P(\pi_i) * \iint P(data \mid \pi_i(\theta_j)) * P(\theta_j)}{\sum_{i=1}^{m} P(\pi_i) * \iint P(data \mid \pi_i(\theta_j)) * P(\theta_j)} \quad \text{(S4)}$$

Analysis with likelihood at optimal parameterization,

$$P(\pi_i(\theta^*) \mid data) = \frac{P(\pi_i(\theta^*)) * P(data \mid \pi_i(\theta^*))}{\sum_{i=1}^{m} P(\pi_i(\theta^*)) * P(data \mid \pi_i(\theta^*))} \quad \text{(S5)}$$

Here 'data' is the behavioral data pooled across the participants for a condition, 'i' is the policy index, θ is {α, c1} for policies 1 and 3-6, and {α} for policy 2, 'j' represents specific parameterization for θ, $\theta^*$ is the optimal parameterization, and m is the number of policies. The integral in equation S4 was over the parameter space and it was a single integral in the case of policy 2.

This way, for each task condition, we had a posterior over the policies indicating the policy that was most likely given the data across the participants. This was done using equation S4. We also had, for each task condition, the optimal parameter values for each policy such that the likelihood of the data across the participants was maximized given the policy at the optimal parameterization. This corresponds to equation S5.

#### 7. Additional objectivity measures

We defined various methods to quantify objectivity at the level of individual participants for each condition. Here are the additional measures which were not described in the main article.

Distortion measure: A measure of how much the externally provided information, like values and probabilities, is distorted when making decisions. We quantify it in the following manner:

$$Distortion\ measure = |2 * \log_{10} c| + \alpha \quad \text{(S6)}$$

This measure is higher when c is farther from '1' (in any direction) and α is more than 0. These parameters have been described in detail section 6 of supplementary methods above. A higher value of distortion may indicate higher subjectivity or lower objectivity.

Non-randomness measure: Absolute value of the slope of the logistic regression, as described in section 6 above.

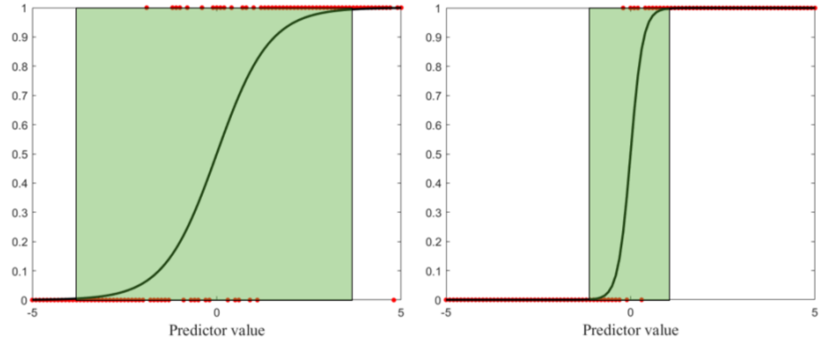

*Figure S2: Illustrative examples to show the interpretation of slope of logistic regression model as indicating uncertainty or randomness in decision policy. Decisions are shown as red dots while the model is shown as a solid black line. The green shaded area shows the part of the model where the probability for selecting either of the options is less than 1. The slope of the model on the left is 1.4 and that on the right is 6.*

A steep (large) slope indicates a policy which has a narrow zone of uncertainty, as shown on the right in figure S2 while a relatively flat (small) slope indicates a policy which has a wide zone of uncertainty, as shown on the left in figure S2 where the decisions relied on 'randomness' or spontaneous subjective influences unaccounted by the policy. A higher value of the non-randomness measure may indicate higher objectivity.

#### 8. Mapping self-reported strategies to candidate policies

Mapping between the participants' responses to the open-ended questions about task-strategy and the policies. If participant's responses were interpreted to mean the statements in column 1, then they were mapped to the policies in column 2. The policy #s correspond to the policies listed in table S3. ST – sure thing value. The notations for gamble characteristics are shown in figure 1B.

*Table S4: Mapping criteria for mapping self-reported task strategy to candidate policies.*

| | **Inferred meaning from the responses** | **Policies to which the response was mapped** |
|---|---|---|
| S1 | Considered points and associated percentages/probabilities | Policies 1, 3, or 4 |
| S2 | S1; Also considered the negative outcome and the associated probability. | Policy 1 |
| S3 | S1; Focused on the highest value and its associated probability | Policy 4 |
| S4 | S1; Focused on the positive outcomes from the gamble and their associated probabilities | Policy 3 |
| S5 | Considered points and associated probabilities of gamble outcomes that were more than ST | Policy 5 |
| S6 | Considered how many gamble outcomes were more than ST | Policy 6 |
| S7 | Considered the points of the gamble outcome that had the highest probability associated with it. | Policy 6 |
| S8 | Considered the probabilities of the gamble outcomes that were more than ST | Policy 2 |

Note that, in strategy S7, the focus is on the gamble outcome associated with the highest probability; this was 'V2' (or the second highest outcome in terms of points in the gamble) for all gambles and hence this strategy is well captured by Policy 6.

#### *9. fMRI acquisition and preprocessing*

The fMRI data was collected using the Echo planar imaging (EPI) in the Siemens 3 Tesla TIM Trio MRI scanner in the Imaging Research Facility at Department of Psychological and Brain Sciences at Indiana University Bloomington. The data was obtained as axial slices at an angle on 30° with anterior-commissure-posterior-commissure line using 64-channel head coil. In a single run lasting 6 mins, 178 volumes of the T2* weighted functional scans were collected with TR = 2000 ms, TE = 25 ms, flip angle = 70°, and 64 x 64 voxel matrix. A single volume consisted of 35 slices of thickness 3.8 mm. One T1-weighted structural scan was collected with TR = 1800 ms, TE = 2.7 ms, flip angle = 9°, and 256 x 256 voxel matrix. This consisted of 160 slices of thickness 1 mm. Preprocessing was performed in SPM12. AFNI's 3dDespike was used for spike correction. The scans were normalized to the standard Montreal Neurological Institute (MNI) space. Spatial smoothing was performed using 8 $mm^3$ full-width-at-half-maximum (FWHM) kernel.

### II. Supplementary Results

#### 1. Bayesian inference of the behavioral policies: Bayes' factor (BF)

This section shows the Bayes' factors of winner policy with respect to other candidate policies (Table S3) where 'winner' policy is the one having highest posterior (Equation S4) given the behavioral data pooled across the participants.

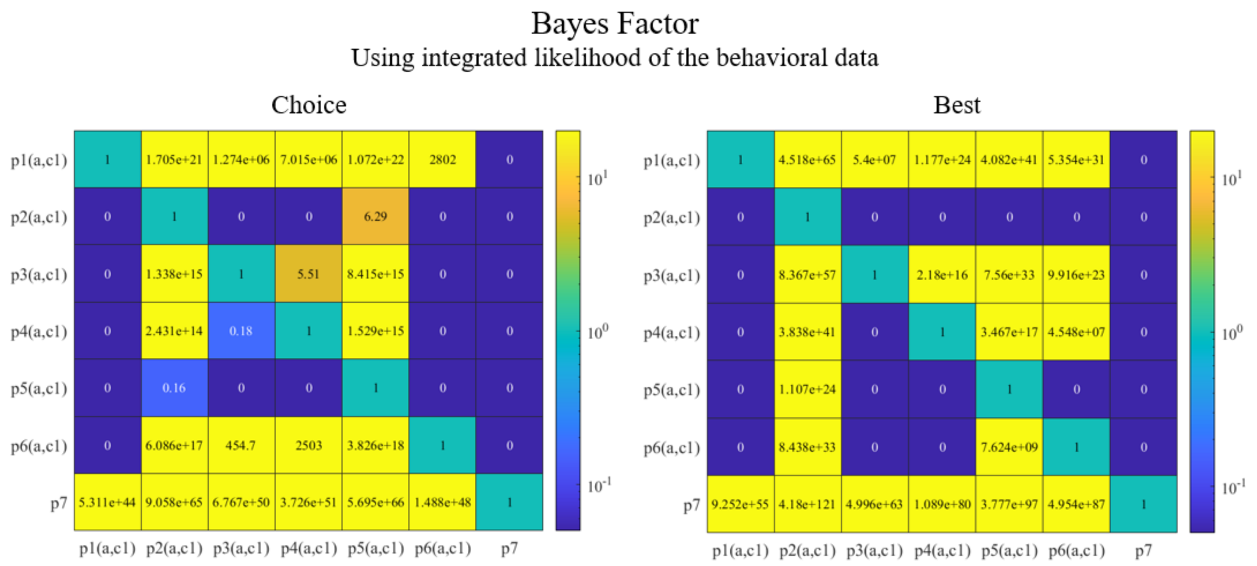

*Figure S3: Bayes Factor, $B_{ij}$, of policy j against policy i where 'i' is the row index and 'j' is the column index for the Choice (left) and Best (right) conditions. The numbers were rounded to two decimal places and hence, any value lower than 0.005 is zero in the figure. A way to utilize colors to quickly read these plots is to see how much yellow does a row have – if a row is all blue, then it means that there is strong evidence for the policy corresponding to that row against all other policies.*

Please see the left panel of figure S3 where each row shows the Bayes' Factor of all policies against the policy representing that row. In row 5, representing policy 5, one can see that BF for all other policies is much lower than 1 except for policy 2. In other words, there is strong evidence for policy 5 against all other policies but policy 2. Even though the BF for policy 2 against policy 5 is more than but still close to 0.1, one cannot say that the evidence for policy 5 over policy 2 is 'strong'. With this caution noted, we take policy 5 to be best describing the behavior in Choice condition moving forward. Please see the right panel of figure S3 for BF of every policy against every other policy in the Best condition. Please note that there is strong evidence for policy 2 against all other policies.

#### 2. Self-reported task strategies in Choice and Best conditions and a summary of the differences between the conditions

Responses to open-ended question about subject's thought process and strategy in the task conditions (column 1 – 'Choice' condition; column 2 – 'Best' condition). Column 3 is the response to what differences there were, if there were any, in these between the choice and best conditions.

*Table S5: Text responses of participants to open-ended questions about task-strategy.*

| Subject # | Choice | Best | Difference between choice and best |
|---|---|---|---|
| **1** | *I chose the gambling one if the number is bigger than the sure answer and have quite decent chance for me to receive it. But sometimes I just want to see if I will win the larger number. If the sure answer is obviously bigger than any other answers, I will choose the sure answer.* | *I choose for the best answer that I am able to have the chance to get the biggest number and receive points.* | *Yes. sometimes I want to try the one that the largest number has the lowest chance because I want to see if I can win the points, but that may not be the best option because the risk of losing point is too high.* |
| **2** | *I tended to want to pick the less risky option but if the possibility of a higher score was likely enough I would go for that. As it went on I got more and more conservative due to loosing a couple of choices.* | *I was more focused on the possibility of a positive value instead of the values themselves.* | *When I was picking the options myself and getting that value I put more thought into the values themselves and my chance of getting more points through each selection. When I was indicating which option was best, it was just about my chances of getting a negative outcome.* |
| **3** | *I chose the best option based on the chances I had to get a higher* | *I chose the best option based on the chances I had to get a higher* | *When I was simply asked to choose I did not have to use my strategy I just had* |

| | | | |
|---|---|---|---|
| | *number on the gambling side or if it was just better to get the for sure points.* | *number on the gambling side or if it was just better to get the for sure points.* | *to look at the diagrams and pick as I was told so it was easier and faster in my head.* |
| **4** | *In the trials in which I chose an option, I considered both the percentages on the graph and the point values associated with them. I preferred the SURETHING option if the more probable option on the GAMBLE graph was lower or close to the point value of the SURETHING option. On the other hand, if there was a potential larger gain (e.g. 30% chance of getting 50 points when the surething option offered 7 points), I would select the GAMBLE option. Generally, decided to select the GAMBLE option if it was a 50% vs 50% and there was more than a point of difference between the positive GAMBLE option and the SURETHING option.* | *For the trials in which I was asked to choose the best option, I made my decision based on what percentage of the GAMBLE graph would result in a higher number of points than the SURETHING graph. I tried to focus on this more than how many more or fewer points were at stake, because I would gain or lose 10 points regardless. Generally, decided to select the GAMBLE option if it was a 50% vs 50%.* | *Yes, I considered the point values more in the trials in which I was asked to choose, but I mostly considered the percentages on the GAMBLE graph that offer more (no matter how many more) points than the SURETHING graph in the trials in which I was asked to indicate the best option.* |
| **5** | *I would first look at the value of the "sure thing" option and then compare that with the values and probabilities of the "gamble" option. Seeing as I only had 5 seconds, to make a decision, I didn't have time to mentally calculate which option would (statistically) grant me the higher value so it was more of a <u>quick mental analyzation</u>. Say* | *I would firstly look at the "sure thing" option and see how many points it was worth. Secondly, I would then look at the "gamble" option and calculate the probability of getting a number greater/less than the sure thing's value. For example, say the "sure thing" option had a value of 20 points with 100% probability and the* | *Yes, there was. When I was asked to indicate the best option, I followed a statistical procedure whereas when I was merely asked to choose, I'd roughly calculate the probability of getting each value (from the gamble option) and would take the safe option (sure thing) if the probabilities weren't favorable or I'd take the "gamble" option* |

| | | | |
|---|---|---|---|
| | *the "sure thing" value was 15 points and the "gamble" option had values of -2, 30, and 40, with respective probabilities of 10%, 40% and 50%, this would be an easy decision to select the "gamble" option as there's only a 10% chance I would lose points and there's a 90% chance I'd gain more points than if I selected the "sure thing" option. However, when the values were mor similar with less favorable probabilities for higher values (and higher probabilities for unfavorable values), I would feel more conflicted and would be more likely to select the "sure thing" option as I was guaranteed the points, regardless how many points I would get.* | *"gamble" option had values of -2, 10, and 30 with probabilities of 20%, 60%, and 20% respectively. I would then calculate that should I select the "gamble" option, the probability of getting a number less than 20 would be 80%. This meant that should I select the "sure thing" option, there was an 80% chance that that would be the best option (and thus grant me 10 points). I used this statistical method for every option. Obviously, I didn't select the best option all the time but I always chose the statistically best option.* | *if the probabilities were favorable.* |
| **6** | *I chose largely based on a risk to benefit analysis. Each one was taken on a case by case basis. To be honest, I couldn't do the math on all of them in my head and so I had to use my intuition on many of them. I usually tried to take risks when I felt the cost was relatively low. I also made decisions based on the opportunity cost. For instance, just because there might only be a 10% chance of losing 8 points if I went with the gamble to gain a higher* | *I used the overall probability of to make my assessment. In other words, if there was one circle with a 100% chance of getting 10 points and another circle with a 30% chance of getting -20 points, a 30% chance of getting 15 points, and a 50% chance of getting 50 points, I would combine the probabilities to determine that there was an 70% chance to for the second option of getting a higher number while there was only a 30%* | *Yes, I chose the options based on how many net points I could be gaining over the course of the game and not on if each individual trial resulted in a higher outcome.* |

| | | | |
|---|---|---|---|
| | *number, there was also the opportunity cost of losing out on the safe point value, such as a 20 point option. Therefore, the risk for a point loss was the negative point value that could be lost, plus the positive point value foregone with the guarantee option.* | *chance of getting a higher number with option one. (i.e. 10 > -20)* | |
| **7** | *For the part that asked me to choose an option, I think it's easier for me to do choose. Because I'm a person who's not good at math and I always think a lot when it comes to making choices (except the things I really want or I need to do). So, I would say there's no strategy in this part, I just followed the instructions without thinking too much as I did in the "best option" part.* | *For the best option, I just think what might be the best choice to win the most point. I didn't really calculate or think that much, I just follow my instinct.* | *For the best option part, I tend to overthink too much even if I just followed my instinct. For the other part, it's just easier to follow instructions and I didn't think that much about how much I'm going to lose. I think it might have some connection with the responsibility feelings in my mind. For instance, for the best option part, I'll tell myself whatever I choose, this is my choice, so I tend to have a responsibility feeling to myself that I need to think properly before choosing. And for another part, I didn't have that much responsibility feeling to myself in my head, I just went for what instruction asked and didn't need to think and worry if the choice was going to affect anything. I'm not sure if this makes senses, but I think it might be one of the reason and difference how I reacted when I did these two parts.* |

| | | | |
|---|---|---|---|
| **8** | *I evaluated both the highest numbers and the probability of me getting those numbers, and my choice based on those two things.* | *The first thing I looked at was the highest number, after which I looked at the probability of me getting that number if I selected that choice and based on that I would choose the gamble or the surething.* | *No, because I utilized the same thought process of evaluating my chances of getting the number I wanted in both tasks in the same manner.* |
| **9** | *When the gamble had about a 70-80% chance of being higher than the surething, I went for the gamble. If it was a 50/50 split of being much higher or much lower than the surething I usually picked based on my previous winning streak (ex: if I lost a lot I went with the surething, but if I had been winning I went with the gamble). If the surething was about the same of the number of majority in the gamble, I just went with the surething.* | *To pick the best option I usually went with surething since it was guaranteed. I picked the gamble only if it was usually 70-80% chance that it would be higher than the surething. If the gamble had a negative more than 20% I didn't choose it.* | *I think I was more careful in the best option compared to the section I was asked to choose. In the section where I was asked to choose, a lot of my decisions depended largely on whether or not I had a lucky or unlucky streak in the options prior.* |
| **10** | *My thought process consisted of comparing the two options, seeing which side had the highest possible benefit while also trying to keep it safe as well. If the gamble numbers were similar to the surething option, or if they were very close, I would always choose the surething option because I was trying to get the most points rather than risk it. If there was a low chance for a higher number, I would more than likely choose the surething option. Being* | *My thought process consisted of just choosing the one that seemed less risky or that could give me more benefit. As time went on in the second section, I realized I could add the percentages of the gamble together if two parts of it were larger than the number in the surething option, so then I would choose the gamble option. Sometimes I got very tired and missed out on an answer because I hadn't realized five seconds had passed.* | *With the best option, I was more likely to choose the less risky option I feel. When I was asked to choose I think that I was more likely to choose the riskier option because of the possible reward; while choosing the best option doesn't always mean riskiest.* |

| | | | |
|---|---|---|---|
| | *able to see what I missed out on by not choosing the other option did affect my strategy, if I saw there was a longer streak of the points being higher in the other option, then I was more likely to choose that option. I would sometimes say "screw it or why not?" and choose the gamble option.* | | |
| **11** | *For the second task, selecting between two options was simple. I would pick the option which indicated more points while using the same strategy described above. Sometimes I would calculate the total percent against SURETHING to make sure I selected the best option. Most of the time, it was about looking at the largest number, and intuition.* | *My strategy for completing this survey or test was simply to look out for the numbers and guess which number is greater. In cases where it indicated that the SURETHING is greater than any other value indicated in the gamble. In cases where the gamble was 50/50, I looked for the number, whether its greater than the SURETHING or not. If yes, I would go for it. In cases where the Gamble showcased the points more than the SURETHING, for example, the numbers are 30 points for 25%, 44 points for 45% and -20 for 30%, and 28 points for SURETHING(100%), I would generally select the Gamble option.* | *NA* |
| **12** | *For questions regarding picking which option was the gamble or the surething, I just picked whichever option had different percentage outcomes as the gamble,* | *For questions regarding picking the best option in terms of point accumulation, I chose what I felt gave me the best chance of earning the most points on every* | *No* |

| | | | |
|---|---|---|---|
| | *and the 100% as the surething.* | *given question. However there were times when I would take gambles and pick an option based on a gut feeling, not rationale thought.* | |
| **13** | *When I was choosing an option, I tended to just chose the higher percentages rather than the point values.* | *When choosing the best option, I mainly focused on how many points I would get. If the point value for the surething graph was higher than the gamble one then I automatically chose that. I only chose the gamble graph if I had a gut feeling or if I saw that the probability of getting more points than sure thing was greater.* | *I took more time in decision making when choosing the best option vs when I was just asked to choose.* |
| **14** | *For the second trial, I chose options based on which majority had the highest amount of points. So if one options majority had more points, I'd usually choose that option.* | *For the first trial, I used the strategy of choosing whichever option I felt had the highest probability of me gaining more points.* | *Yes, for the ones in which I was asked to choose the best option, I usually choose whichever option I believed had the best probability and security of me getting points. For when I was just choosing whichever option, I went with whenever the highest probability if it had more points than the other options.* |
| **15** | *At first, I picked the "safest" option, meaning I only picked gamble if the option for losing points was 20% or less. This is the same strategy I used for the part 1 remote section. But over time, I began reverting to my strategy from task 1, meaning only picking gamble if the largest sector had a point value* | *Over time I developed a strategy; if the surething point value was greater than the largest sector percentage point value in gamble, I would pick surething. But if the largest sector in gamble had a point value greater than surething, I would pick gamble.* | *When I was asked to choose, I played it a little safer, because even if I knew the "best" option would allow me to net greater points received over time, I don't like to see large amounts of points lost, or any at all. So, I played it more safe with task 2 at first, but over time used the same* |

| | | | |
|---|---|---|---|
| | *greater than surething. This seemed to get me better results.* | | *strategy as task 1 with the best option.* |
| **16** | *I first looked at the risk of the situation—that is, how much I could lose in the Gambling circle. If it was 20 or over, I was very unlikely to choose it unless the Sure thing reward was very low and the Gambling reward very high. If it was an intermediate risk such as 10, I was likely to calculate the probability of the Gambling reward and make a judgment call based on the value of the Sure thing circle.* | *I would calculate the likeliest outcome for the Gambling circle and compare its value to the Sure thing circle. I would also consider the second likeliest outcome for the Gambling circle. If it was a worse outcome than the Sure thing circle and its probability combined with the "negative" section of the Gambling circle was 35-40%, I was likelier to choose the Sure thing circle.* | *Yes, I was much more concerned with possible risk for the "ask to choose" task. I was likelier to consider strict probability for the "best option". If the risk was lower, I was likelier to take chances for rewards for the "ask to choose" tasks.* |
| **17** | *As described above, I took an approach that I felt would maximize my wins while minimizing my losses. So I would look at the available point values and make my choices based on that. The percent chance of getting the different point values was less of a concern. So if I had a 100% chance of getting 30 points, compared to a 50% chance of 25 points and a 20% chance of 60 points, I would choose the 30 points. My evaluation changed depending on the point value. It felt like if the surething choice was 50% or less than the gamble options, I would choose the gamble options because that would maximize my* | *At first, I was treating this task as I treated the similar task in part 1 (or the second set of tasks in part 2). That is, I was choosing the option that I felt would most likely maximize my wins while minimizing my loses. After many rounds in a row where I lost points because the option I chose resulted in a slightly lower value than the alternative, I changed my tactic. I started thinking a little more risky and choosing the option with the highest possible point outcome. For example, if the surething choice was 30 points, but the gamble choice presented the option of 50% chance of 20 points, 20% chance of 40 points, and 30%* | *Yes, as I described above. When I was asked to choose the best option with no regard for total points gained/lost, I was more confident in risky-er choices. When I was choosing between different real point values, my choices were more measured.* |

| | | | |
|---|---|---|---|
| | *winning potential. Interestingly, I did not really factor in the negative point values. I even found myself surprised once that I chose the gamble option when there was a 30% chance of loosing 20 points.* | *chance od -10 points, I would chose the gamble option. I feel like I had more success choosing risky options in this task.* | |
| **18** | *I paid attention to what they were asking but it was a bit hard because I was getting tired. I was also not as quick because sometimes the sure thing would be on the right for like 2 or 3 turns then it would switch to the other side.* | *In the beginning I was just trying to use the percentages as a way to help. I was thinking like which one would be greater and should I gamble and try to get more points or choose the sure thing. Towards the middle I was just trying my best to do what the task asked me to do but I was getting tired, so I tried my best to be quick but still be as accurate as I can in my answers. In one of the tasks towards the end, I tried to use the colors of the circles to pick my answers. Like if the sure thing was yellow, I would look and see how much points the yellow piece of the other circle was. If it was greater than the sure thing I would gamble or choose the gamble instead of the sure thing.* | *No I used the same strategies that I put down for question 1.* |
| **19** | *I first read the sure thing option. Then I read the percentages of each value on the gamble option. The gamble option normally had a 50% or 40% value then a 20-30% value then* | *I read the number for the sure thing option then I read the 2 highest numbers from the gamble option. Normally the gamble option was 70% or 80% >0. On the* | *Yes, there was a difference. During the best option, I just looked at the 2 highest values on the gamble option, and if they were both higher than the sure thing value,* |

| | | | |
|---|---|---|---|
| | *another 20-30% value that is negative. If the negative value was small AND the other 50% value was within a couple points of the sure thing option, I would choose the gamble option. If the negative value was high and the other 50% value was way lower than the sure thing option, I would choose the sure thing option.* | *gamble option, if there were 2 options that were higher than the sure thing option, I would choose the gamble option. If there was only 1 option that was higher than the sure thing option, I would choose the sure thing option.* | *I would choose the gamble option. And if one of the values were lower than the gamble option, I would choose the for sure thing option because I would only have a 20-30% chance of loosing points. During the choice option, I actually had to pay attention to the neative points and how close the high percentage value was to the sure thing value. If it was close than there was a 50% change of me getting within 5 points, a 20% chance of me loosing a SMALL negative value, and a 20-30% chance of me getting a large value of points.* |
| **20** | *I tried to calculate the expected value by multiplying the numbers by the percentage and adding them up and comparing it to the 100% guaranteed option, though it was a bit hard to do in 5 seconds so I had to guess for most of them.* | *I tried to see if the value of the guaranteed outcome had a lower probability of occurring compared to the other one (for example if it was 9.98 guarenteed, and the other choice was 50% 10, 20% -10, and 30% 30, I realized that the second one was 80% more likely at winning.* | *For the best option, it was more based on just getting a better outcome so it was much easier compared to choosing because I had to try to maximize my points outcome.* |
| **21** | *The strategy I used again was the basic probability with a margin for better profits. 1. If the 100% option yielded better results when the gable profits combined, then I chose surething. 2. If the value of maximum probability of the gamble option is greater the surething value then I* | *The strategy I used was the basic probability. 1. If the 100% option yielded better results when the gable profits combined, then I chose surething. 2. If the value of maximum probability of the gamble option is greater than the sure thing value then I chose gamble.* | *Yes, in case of the choosing the best option, the points I could gain was the same as the points I would lose. Therefore I really didn't have to consider the gamble option more seriously unless there is a clear winner in the charts. But in case of choosing the option, there is a different* |

| | | | |
|---|---|---|---|
| | *chose gamble. 3. If the probability of maximum value was minimum, however the delta of the sure thing and the maximum profit was more then I chose to go with the gamble. 4.If the probability of maximum value was minimum, but the loss delta is more than the profit margin, I chose the surething option.* | | *in points I could gain, therefore I changed the way I considered the gamble options, based on the maximum I could gain.* |
| **22** | *I think I strategies by for gambling, adding the percentage of the positive points and comparing it to the negative. Or just looking at how small or large the difference is between the sure thing option and gambling option. Sometimes 50% of gamble was similar to the sure thing option, so I hesitated a little but most of the time, I was able to pick the one that had more points.* | *For choosing the best option, I was adding the percentage if they had greater value than the sure thing option. But also if the gamble option had too much of a negative points for the percentage, I would choose the sure thing option. I was just going with the flow and seeing if the gamble would give me a better option, but some of the question were easy to determine than others.* | *Best option was where I choose based off of the percentage more than the points that were on the screen. If it had more percentage of winning on the gamble, I would try to go for the gamble instead of the sure thing option. For accumulating points, I was looking more at the points and seeing the odds for me to get more points, which when I come to think about it, it is a same thought process but more percentage for the best option questions.* |
| **23** | *The thought process for this experiment consisted more of looking at the probabilities of the gambling option and tallying them up to see if it was more likely than the sure thing option. I thought this experiment felt easier however the 50/50 gambling option was very frustrating.* | *With this experiment, it was very hard to react quickly and see the different numbers and percentages on the gambling option. With the 5 second time constraint, it felt like it was more rushed compared to other experiments. My strategy for this one was trying to see if the surething option had a better chance of landing than the* | *I'd say my thought process for the first one was slightly different compared to the second one. I felt like the first one was more rushed as I was trying to take in all the options and choose either gambling or the sure thing option. The second experiment I would look at the sure-thing immediately and then see if it had a higher chance compared to the gambling* |

| | | | |
|---|---|---|---|
| | | *gambling option and its percentages.* | *option or vise versa. Overall, I thought the first one was much more reliant on reaction time.* |
| **24** | *No strategy* | *No strategy, just fillings* | *When I were asked to indicate the best option, I tried to choose the least risky one* |
| **25** | *When I was asked to indicate the best option, I spent less time considering the gambling option and quickly realized that the surething option was more reliable and better in the long run.* | *When I was asked to indicate the best option, I spent more time considering the gambling option in case it reaped better rewards with little risk.* | *Yes; when I was asked to indicate the best option, the pros and cons of each option was less clear. But once I was asked to choose, it was clear that surething option was better.* |
| **26** | *First I looked at the point values associated with the options. I naturally wanted to go with the higher point value but needed to evaluate the chances. If the "sure thing" option had a point value similar to the highest point value for the "gamble" I would choose the "sure thing". I also took the approach where if the gamble had a combined chance of >70% to get more points than the "sure thing" then I would go for the gamble.* | *To indicate the best option, I took the same approach as described in number two, however I was less risky with my decisions. I tried to objectively look at both options to determine which was better, which was difficult as there was very little time, and I was swayed by point value. I told myself if I was unsure then it would be safer to go with the "sure thing" option.* | *When I was asked to choose, I took more risks depending on how I thought I was doing. I also went off of my gut feeling. When asked to choose the best option I think I may have thought too hard about it, as I was trying to be less risky, which I believe I ended up doing worse at.* |
| **27** | *The first section I was utilizing the percentages as well as the gains (i.e 30% chance for 64 vs a 60% of 15, I would choose this option if the SURETHING was below 15 points but if it was above i.e. 16.01points I would not choose the GAMBLE). I very rarely chose looked at how many* | *This section I focused on collecting all of he GAMBLE percentages together and compared it to the SURETHING. I did not look towards the amount of points per say but rather if those points were above/below the SURETHING plus their percentages (i.e. SURETHING was 15.01* | *As I mentioned above, the choose section felt like I had more freedom, the risk vs reward stimuli was much more engaging and when I took a risky GAMBLE I felt the consequences or reaped the rewards. It felt fairer. However, the best option felt like it was less fair. I utilized adding the* |

| | | | |
|---|---|---|---|
| | *points I would be liable to lose (i.e. 10% to lose 10 points). This section I felt the most free on, it felt the most like a gamble since the risks and the rewards had tangible correlations that I could grasp (i.e. I felt it when I gained 64 points over 15 or lost 20 points etc.).* | *points and the GAMBLE was 60% 14 20% 60 and 20% -7, I would most likely choose the SURETHING as the GAMBLE had an 80% chance of failure.) This section, oddly enough, felt the least fair. The amount I was gambling did not feel tangible so when I would choose the "safe" (according to me) option and still lose it felt extra bad since there really wasn't a concept for "risk and reward" in this section. I felt that I had less freedom compared to task 1.* | *percentages together over looking at the amount of points I could win. I felt like I had less freedom because what I was gambling was not exactly up to my discretion.* |
| **28** | *I only chose the gambling option if there was a 50% chance or greater that I would get a higher value than the sure thing value.* | *For these trials I thought about combining any section that had a greater value than the sure thing value and if those sections were more than 50% then I chose that option.* | *The choose trials were a little different for me because I might have still chosen to gamble with an amount that was only slightly less than the sure thing value. For example, maybe the sure thing value was 20. If the gambling values were 45% for 30 and 20% for 18, I might have still picked the gambling option because the 18 value was not that much less than the 20 value. This was different in the best trials, though, as I only picked the gambling option if there was a 50% chance or greater of me getting a higher value than the sure thing value.* |
| **29** | *I used the same strategy as above.* | *I kept a number from one of the options which was 100% to consider and* | *No, there wasn't any difference in my thought process, I was using the* |

| | | | |
|---|---|---|---|
| | | *then assumed the probability from the second option which said 50% for some number, 30% for some number and 20%, it was something like that. So then again, I calculated which option weighs more, then I finalized that option.* | *same strategy at all times i.e I always selected the best option.* |
| **30** | *I chose the sure thing if the number was high enough and the gamble probabilities weren't too bad, however if the gamble probabilities were worth it if they gave out a significant higher number compare to the sure thing, then I'd risk it.* | *I chose what I thought was the best option pretty much. Only other thought process behind it was taking a minor risk by gambling.* | *Choosing (task 1) was easier since there wasn't any risk in losing points if the outcome was lower then the other choice. Task 2 require some thought since I was risking losing points if the result was lower.* |

<u>Summary of the differences reported between both the conditions:</u>

- 8 participants mentioned that they focused on specific outcome values more in Choice than in Best. 1 participant mentioned the opposite.
- 8 participants mentioned making decisions based on more intuition/gut-feeling/rough-calculations in Choice than in Best. 2 participants mentioned the opposite.
- 6 participants reported finding the Choice task less difficult to perform or leading to less tiredness than the Best. 1 participant reported the opposite.
- 5 participants mentioned more risk-seeking and/or exploration in choice than in the best. 2 participants mentioned the opposite.
- 2 participants reported more learning from task experience in Choice than in the Best.
- 1 participant reported taking less time to decide in Choice than in the Best.
- 1 participant reported thinking more about the long-term rewards from options in Choice than in the Best.

Please note that more participants may have experienced these (or opposite) differences between the tasks but may have simply not mentioned so in their responses to the open-ended questions. The numbers above only account for participants that explicitly mentioned these factors.

#### 3. Behaviorally inferred and self-reported policies of individual participants

We performed model comparison at the level of individual participants' data as well as collected self-report of their task strategies. The self-report was mapped onto the candidate policies, with mapping shown in supplementary methods section 8. This section shows the behaviorally inferred policies and the policies that the self-report was mapped to.

*Table S6: Policies that the responses on task-strategy are mapped to (left) and policies inferred from model comparison based on behavioral data (right).*

| ***Participant*** | | **Policies from self-report** | **Policies from behavioral data** |
|---|---|---|---|
| ***1*** | Choice | 2, 5, 6 | 2 ($\alpha$ = 0.21) |
| | Best | NA | 3 (c1 = 0.46) |
| ***2*** | Choice | 5 | 4 (c1 = 0.73), 5 (c1 = 0.73) |
| | Best | 2 | 3 (c1 = 0.65) |
| ***3*** | Choice | NA | 4 (c1 = 0.54), 5 (c1 = 0.54) |
| | Best | NA | 4 (c1 = 0.70, $\alpha$ = 0.12), 5 (c1 = 0.70, $\alpha$ = 0.12) |
| ***4*** | Choice | 5 | 6 (c1 = 0.61) |
| | Best | 2 | 5 (c1 = 0.84) |
| ***5*** | Choice | 1, 2 | 3 (c1 = 0.64) |
| | Best | 2 | 2 |
| ***6*** | Choice | 1 | 1 (c1 = 0.85) |
| | Best | 2 | 5 (c1 = 0.90, $\alpha$ = 0.06) |
| *7* | Choice | NA | 3 |
| | Best | NA | 3 |
| ***8*** | Choice | 3,4 | 6 |
| | Best | 4 | 2 |
| ***9*** | Choice | 2 | 6 |
| | Best | 2 | 2 |
| ***10*** | Choice | 3, 4 | 5 (c1 = 0.57) |
| | Best | 2 | 2 |
| ***11*** | Choice | 2, 4 | 6 (c1 = 1.49) |
| | Best | 2 | 5 (c1 = 0.61, $\alpha$ = 0.05) |
| ***12*** | Choice | NA | 5 (c1 = 0.44) |
| | Best | NA | 6 (c1 = 0.56) |
| ***13*** | Choice | 2 | 6 |
| | Best | 1, 3, 4 | 2 |
| ***14*** | Choice | 6 | 5 (c1 = 1.58, $\alpha$ = 0.55) |
| | Best | 2 | 3 (c1 = 1.32, $\alpha$ = 0.22) |
| ***15*** | Choice | 1, 6 | 2 |
| | Best | 6 | 6 |

| | | | |
|---|---|---|---|
| ***16*** | Choice | 1 | 1 (c1 = 0.82) |
| | Best | 2 | 4 (c1 = 1.2) |
| ***17*** | Choice | 3 | 5 (c1 = 0.76, α = 0.15) |
| | Best | 4, 5 | 2 |
| ***18*** | Choice | NA | 1 |
| | Best | 2 | 1 (c1 = 0.64) |
| ***19*** | Choice | 1 | 4 (c1 = 0.65) |
| | Best | 6 | 2 |
| ***20*** | Choice | 1 | 2 |
| | Best | 2 | 2 |
| ***21*** | Choice | 2, 4, 6 | 2 |
| | Best | 6 | 2 |
| ***22*** | Choice | 1, 2 | 5 (c1 = 0.46, α = 0.09) |
| | Best | 2 | 2 |
| ***23*** | Choice | 2 | 1 |
| | Best | 2 | 2 |
| ***24*** | Choice | NA | 5 (c1 = 1.34) |
| | Best | NA | 2 (α = 0.58) |
| ***25*** | Choice | NA | 3 |
| | Best | NA | 3 (c1 = 1.3, α = 0.67) |
| ***26*** | Choice | 2 | 3 |
| | Best | 2 | 4 (c1 = 0.72, α = 0.30) |
| ***27*** | Choice | 3 | 6 (c1 = 0.75) |
| | Best | 2 | 6 (c1 = 0.85) |
| ***28*** | Choice | 2 | 6 (c1 = 0.92) |
| | Best | 2 | 2 (α = 0.000036) |
| ***29*** | Choice | NA | 6 (c1 = 1.82) |
| | Best | NA | 6 |
| ***30*** | Choice | NA | 3 (c1 = 0.79) |
| | Best | NA | 5 (c1 = 0.32) |

4. Objectivity measures: Obtained from behavioral data pooled across the participants for each condition

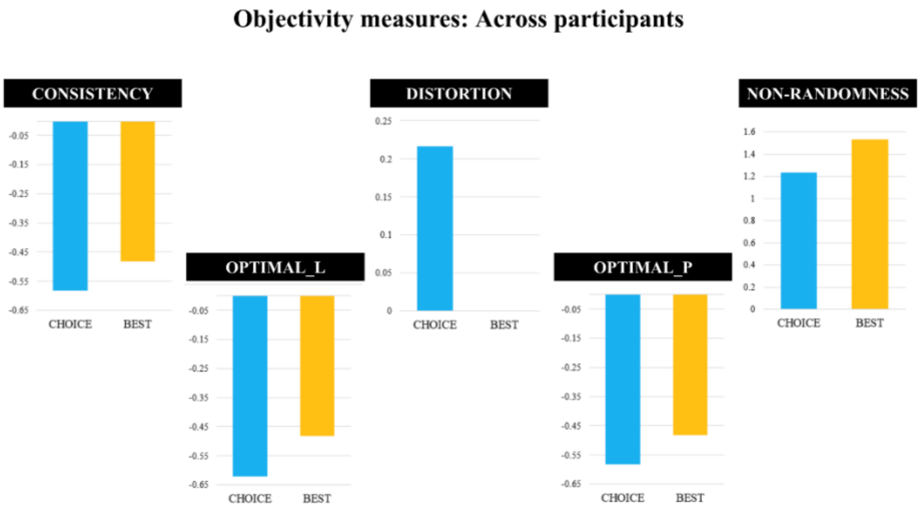

*Figure S4: Objectivity measures obtained from analysis of behavioral data pooled across participants.*

#### 5. Order effects in Choice – Best contrast: Whole brain exploratory analysis

We conducted a whole-brain exploratory two-sample t-test for the main and the ($Decision_{Choice}$ – $Decision_{Best}$) contrast. For order 1 > order 2, no cluster reached significance. Table S7 and figure S5 show the results for order 2 > order 1.

*Table S7: Results for whole-brain exploratory two-sample t-test comparing groups 1 and 2 representing different orders in which the Choice and the Best blocks were administered.*

| | | | | **Peak MNI coordinates** | | | | |
|---|---|---|---|---|---|---|---|---|
| | Region | **Laterality** | **Cluster Size** | **X** | **Y** | **Z** | **Max stat t** | **P Cluster Corrected** |
| **1.** | **($Decision_{Choice}$ – $Decision_{Choice\_control}$) – ($Decision_{Best}$ – $Decision_{Best\_control}$): Group 2 > Group 1** | | | | | | | |
| | Posterior cingulate, BA 29, BA 30 | Bilateral | 212 | 8 | -48 | 14 | 5.25 | 0.04 (FDR), 0.061 (FWE) |
| | | | | -10 | -56 | 10 | | |
| **2.** | **$Decision_{Choice}$ – $Decision_{Best}$: Group 2 > Group 1** | | | | | | | |
| | Posterior cingulate, Cuneus, Cingulate gyrus, BA 31 | Bilateral | 1868 | 6 | -58 | 10 | 7.02 | <0.001 |
| | | | | -2 | -62 | 12 | | |

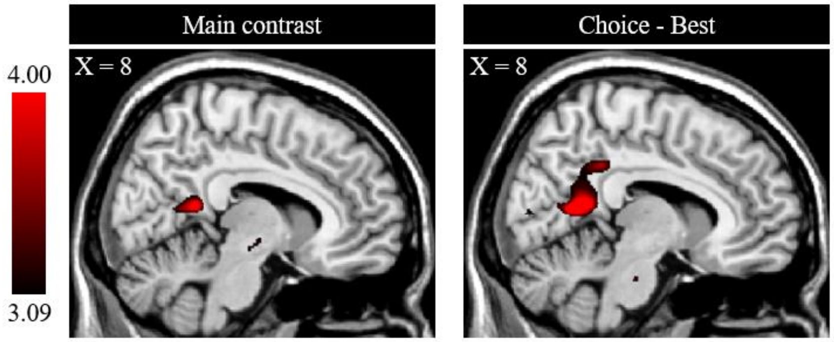

*Figure S5: Whole-brain exploratory two-sample t-test across the participants comparing groups with different sequences of task-block, Group 2 > Group 1. Clusters shown have significantly greater effect of the contrasts for group 2 than group 1. Group 1 corresponds to order 1, that is Choice → Best, and group 2 corresponds to order 2, that is, Best → Choice.*

#### 6. Additional fMRI results' tables

'C', 'B', and 'Cntrl' refer to the contrast $Decision_{Choice}$, $Decision_{Best}$, and $Decision_{Control}$ respectively. Contrast*x represents results of parametric modulation of the contrast with x. Unless specified otherwise, the results correspond to whole brain exploratory analysis in SPM with cluster defining threshold equal to 0.001.

**C – B**

| | | | | Peak MNI coordinates Clust thresh = 0.008 | | | |
|---|---|---|---|---|---|---|---|
| **Region** | **Laterality** | **Cluster Size** | **X** | **Y** | **Z** | **Max stat t** | **P Cluster Corrected** |
| Cingulate gyrus, Mid-cingulate | Bilateral | 1029 | -10 | -36 | 42 | 5.02 | 0.013 |
| | | | 6 | -18 | 58 | | |

**B – C**

No significant clusters

**C – CntrlC minus B – CntrlB**

| | | | | Peak MNI coordinates | | | |
|---|---|---|---|---|---|---|---|
| **Region** | **Laterality** | **Cluster Size** | **X** | **Y** | **Z** | **Max stat t** | **P Cluster Corrected** |

| | | | | | | | |
|---|---|---|---|---|---|---|---|
| Cingulate gyrus, Mid-cingulate, middle frontal gyrus, BA 24, BA 31 | Bilateral | 328 | -8 | -18 | 48 | 4.57 | 0.011 |
| | | | 6 | -18 | 58 | | |

**B – CntrlB minus C – CntrlC**

No significant clusters.

**C – CntrlC**

| | | | Peak MNI coordinates | | | | |
|---|---|---|---|---|---|---|---|
| **Region** | **Laterality** | **Cluster Size** | **X** | **Y** | **Z** | **Max stat t** | **P Cluster Corrected** |
| Precuneus / Parietal lobe / BA 7, Mid occipital lobe | Bilateral | 8693 | -10 | -66 | 52 | 12.02 | <0.001 |
| | | | 10 | -62 | 52 | | |
| Thalamus / Medial dorsal nucleus, Lentiform nucleus / Medial globus pallidus | Bilateral | 2710 | 8 | -18 | 10 | 8.88 | <0.001 |
| | | | -8 | 0 | -4 | | |
| Cingulate gyrus / mid-cingulum, Anterior cingulate / BA 32, Supplementary motor area, Frontal superior gyrus, Superior frontal gyrus | Bilateral | 3826 | 2 | 22 | 34 | 8.83 | <0.001 |
| | | | -24 | -2 | 52 | | |
| Inferior frontal operculum, Inferior frontal gyrus / BA 46, | Right | 1423 | 40 | 4 | 24 | 6.99 | <0.001 |

| | | | | | | | |
|---|---|---|---|---|---|---|---|
| Middle frontal gyrus | | | | | | | |
| Insula / Inferior frontal gyrus | Right | 272 | 32 | 20 | -6 | 6.63 | 0.026 |
| Inferior frontal gyrus / BA 9, Precentral gyrus | Left | 880 | -46 | 4 | 30 | 6.49 | <0.001 |
| Declive (Cerebellum), Culmen/Vermis_6 | Bilateral | 668 | -6 | -74 | -26 | 6.47 | <0.001 |
| | | | 4 | -58 | -24 | | |
| Declive (Cerebellum), Culmen | Left | 665 | -36 | -58 | -30 | 6.32 | <0.001 |
| Fusiform gyrus | Right | 298 | 44 | -58 | -18 | 5.18 | 0.018 |
| Middle occipital gyrus | Left | 193 | -38 | -92 | 2 | 4.82 | 0.086 (FWE), 0.023(FDR) |

**<u>CntrlC – C</u>**

| | | | **Peak MNI coordinates** | | | | |
|---|---|---|---|---|---|---|---|
| **Region** | **Laterality** | **Cluster Size** | **X** | **Y** | **Z** | **Max stat t** | **P Cluster Corrected** |
| Inferior parietal lobule / BA 40, supramarginal gyrus, Insula / BA 13, superior temporal gyrus / BA 41, middle temporal gyrus, postcentral gyrus, precentral gyrus, BA 21, BA 22, BA 42, Uncus / BA 28, Parahippocampal gyrus, Amygdala, Hippocampus | Right | 5669 | 52 | -36 | 26 | 8.28 | <0.001 |
| | | | 24 | 8 | -24 | | |
| Middle temporal gyrus / BA 21, Superior | Left | 7758 | -66 | -38 | -4 | 7.92 | <0.001 |

| | | | | | | | |
|---|---|---|---|---|---|---|---|
| temporal gyrus, Parahippocampal gyrus, Hippocampus, BA 28, Insula, BA 13, Inferior frontal gyrus / BA 47, Amygdala | | | | | | | |
| | | | -24 | -10 | -20 | | |
| | | | -60 | -48 | 42 | | |
| Cerebellum posterior lobe, Cerebellum Crus 1, Cerebellum Crus 2 | Left | 295 | -26 | -92 | -28 | 7.34 | 0.019 |
| Precuneus / BA 31, Cingulate gyrus, Mid-cingulum, Posterior cingulate / Retrosplenial cortex | Left | 440 | -10 | -48 | 34 | 6.98 | 0.003 |
| Cerebellum posterior lobe, Cerebellum Crus 1, Cerebellum Crus 2 | Right | 787 | 32 | -84 | -40 | 6.81 | <0.001 |
| BA 11, Middle frontal gyrus, Orbito-frontal gyrus, Anterior cingulate, BA 32, BA 10, BA 9, BA 25 | Bilateral | 2490 | -6 | 32 | -12 | 6.66 | <0.001 |
| | | | 6 | 52 | 4 | | |
| Mid-cingulum, Cingulate gyrus | Bilateral | 315 | 8 | -18 | 38 | 5.35 | 0.014 |
| | | | -10 | -18 | 38 | | |
| Occipital lobe, Cuneus, BA 18 | Left | 222 | -8 | -92 | 4 | 4.69 | 0.055 (FWE), 0.022 (FDR) |

**<u>B – CntrlB</u>**

| | | | Peak MNI coordinates | | | | |
|---|---|---|---|---|---|---|---|
| **Region** | **Laterality** | **Cluster Size** | **X** | **Y** | **Z** | **Max stat t** | **P Cluster Corrected** |
| Superior parietal lobule / BA 7, Precuneus / BA 7 / BA 39, Middle occipital gyrus | Bilateral | 8629 | 10 | -66 | 52 | 12.93 | <0.001 |
| | | | -10 | -68 | 56 | | |
| Thalamus, Medial global pallidus, Lentiform nucleus, Medial dorsal nucleus, Caudate, mid-brain, Bleeds into anterior cerebellum, Pallidum | Bilateral | 4026 | 10 | 6 | 0 | 8.28 | <0.001 |
| | | | -8 | 2 | -2 | | |
| Cingulate gyrus / Mid cingulum, BA 32, Middle frontal gyrus / BA 8, Supplementary motor area | Right | 1277 | 2 | 18 | 40 | 8.14 | <0.001 |
| Medial occipital gyrus / BA 18, Fusiform gyrus, Cerebellum_Crus1_L, | Left | 1731 | -44 | -58 | -18 | 7.54 | <0.001 |
| Insula, Inferior frontal gyrus / BA 47 | Left | 215 | -30 | 20 | -4 | 7.52 | 0.064 (FWE), 0.011 (FDR) |
| Precentral gyrus / BA 6, Inferior frontal gyrus | Left | 413 | -42 | 0 | 28 | 6.95 | 0.004 |
| Middle frontal gyrus / BA 6, Precentral gyrus | Right | 659 | 34 | 0 | 56 | 6.87 | <0.001 |
| Insula, Inferior frontal gyrus / BA 47 | Right | 344 | 34 | 18 | -4 | 6.77 | 0.01 |
| Cerebellum_9, Cerebellum, posterior lobe | Right | 173 | 10 | -60 | -50 | 6.61 | 0.123 (FWE), 0.02 (FDR) |
| Cerebelum_Crus1_L, Cerebellum posterior lobe, Vermis | Bilateral | 452 | 4 | -76 | -24 | 6.49 | 0.003 |
| | | | -6 | -74 | -28 | | |

| | | | | | | | |
|---|---|---|---|---|---|---|---|
| Middle frontal gyrus, Precentral gyrus | Left | 380 | -28 | -4 | 54 | 6.16 | 0.006 |
| Middle frontal gyrus / BA 46, Inferior frontal gyrus, | Right | 1236 | 48 | 10 | 24 | 5.85 | <0.001 |
| Cerebellum_9, Cerebellum anterior lobe, Vermis, Cerebellum posterior lobe | | 273 | 0 | -58 | -38 | 5.36 | 0.027 |
| | | | -12 | -52 | -44 | | |

**<u>CntrlB – B</u>**

| | | | Peak MNI coordinates | | | | |
|---|---|---|---|---|---|---|---|
| **Region** | **Laterality** | **Cluster Size** | **X** | **Y** | **Z** | **Max stat t** | **P Cluster Corrected** |
| Cingulate gyrus / BA 31, Mid cingulum, Precuneus, Superior temporal gyrus / BA 39, Middle temporal gyrus (Bleeds into Inferior and superior temporal gyrus / BA 21), Supramarginal gyrus / BA 40, Inferior parietal lobule, | Bilateral | 19385 | 64 | -38 | 30 | 8.21 | <0.001 |
| | | | -52 | -8 | -26 | | |
| Mid orbitofrontal gyrus, Anterior cingulum, Inferior frontal gyrus, Superior frontal gyrus / BA 10, Middle frontal gyrus / BA 10 | Bilateral | 2934 | -4 | 38 | -12 | 6.11 | <0.001 |
| | | | 8 | 26 | -18 | | |

| | | | | | | | |
|---|---|---|---|---|---|---|---|
| Hippocampus, Parahippocampus | Left | 451 | -30 | -18 | -22 | 5.46 | 0.003 |
| Superior occipital lobe, Cuneus, BA 18 | Right | 544 | 14 | -82 | 26 | 5.41 | 0.001 |
| Superior occipital lobe, Cuneus, BA 18, BA 19 | Left | 253 | -12 | -90 | 22 | 5.07 | 0.036 |
| Sub-gyral, Caudate | Right | 161 | 24 | 8 | 20 | 4.10 | 0.149 (FWE), 0.056 (FDR) |

*Parametric modulation: ST*

**(C – B)*ST**

No significant clusters.

**(B – C)*ST**

No significant clusters.

**(C – CntrlC minus B – CntrlB)*ST**

No significant clusters.

**(B – CntrlB minus C – CntrlC)*ST**

| | | | | **Peak MNI coordinates (Clust def thresh = 0.005)** | | | |
|---|---|---|---|---|---|---|---|
| **Region** | **Laterality** | **Cluster Size** | **X** | **Y** | **Z** | **Max stat t** | **P Cluster Corrected** |
| Middle occipital gyrus / BA 18 | Left | 947 | -2 | -104 | 10 | 4.28 | 0.003 |

**C*ST**

| | | | Peak MNI coordinates (Clust def thresh = 0.005) | | | | |
|---|---|---|---|---|---|---|---|
| **Region** | **Laterality** | **Cluster Size** | **X** | **Y** | **Z** | **Max stat t** | **P Cluster Corrected** |
| Precentral gyrus / BA 43, bleeds into superior temporal gyrus | Right | 971 | 56 | -8 | 8 | 5.33 | 0.004 |

**Minus C*ST**

| | | | Peak MNI coordinates (Clust def thresh = 0.005) | | | | |
|---|---|---|---|---|---|---|---|
| **Region** | **Laterality** | **Cluster Size** | **X** | **Y** | **Z** | **Max stat t** | **P Cluster Corrected** |
| Cerebellar_9, Cerebellar tonsil, Cerebellum posterior lobe | Bilateral | 1890 | -4 | -52 | -46 | 5.24 | <0.001 |

**(C – CntrlC)*ST**

No significant clusters.

**(CntrlC – C)*ST**

| | | | Peak MNI coordinates (Clust def thresh = 0.005) | | | | |
|---|---|---|---|---|---|---|---|
| **Region** | **Laterality** | **Cluster Size** | **X** | **Y** | **Z** | **Max stat t** | **P Cluster Corrected** |
| Precuneus | Right | 574 | 22 | -60 | 26 | 5.24 | 0.046 |

**B*ST at the time of decision**

No significant clusters.

**Minus B*ST**

| Region | Laterality | Cluster Size | Peak MNI coordinates X | Y | Z | Max stat t | P Cluster Corrected |
|---|---|---|---|---|---|---|---|
| BA 7, Precuneus, Parietal lobe, bleeds into Occipital lobe | Left | 514 | -20 | -68 | 28 | 5.25 | 0.001 |
| BA 7, Precuneus, Parietal lobe, bleeds into Occipital lobe | Right | 450 | 22 | -66 | 26 | 4.95 | 0.002 |

**(B – CntrlB)*ST**

No significant clusters.

**(CntrlB – B)*ST**

No significant clusters.

*Parametric modulation: V2*

**(C – B)*V2**

| Region | Laterality | Cluster Size | Peak MNI coordinates X | Y | Z | Max stat t | P Cluster Corrected |
|---|---|---|---|---|---|---|---|
| Lingual gyrus, Parahippocampal gyrus, BA 19 | Right | 196 | 26 | -56 | -6 | 5.48 | 0.072 (FWE), 0.078 (FDR) |

**(B – C)*V2**

| | | | Peak MNI coordinates | | | | |
|---|---|---|---|---|---|---|---|
| **Region** | **Laterality** | **Cluster Size** | **X** | **Y** | **Z** | **Max stat t** | **P Cluster Corrected** |
| Postcentral gyrus, BA 1, BA 2 | Left | 218 | -48 | -28 | 56 | 5.54 | 0.05 (FWE), 0.04 (FDR) |

**(C – CntrlC minus B – CntrlB)*V2**

No significant clusters.

**(B – CntrlB minus C – CntrlC)*V2**

No significant clusters.

**C*V2**

| | | | Peak MNI coordinates | | | | |
|---|---|---|---|---|---|---|---|
| **Region** | **Laterality** | **Cluster Size** | **X** | **Y** | **Z** | **Max stat t** | **P Cluster Corrected** |
| Superior temporal gyrus, BA 22, Inferior parietal lobule, BA 40, Supramarginal gyrus, Middle temporal gyrus, BA 22 | Right | 991 | 66 | -52 | 12 | 7.10 | <0.001 |
| Anterior cingulate (BA 32), Middle frontal gyrus (BA 9) | Bilateral | 993 | -4 | 48 | 6 | 6.50 | <0.001 |
| | | | 6 | 46 | 20 | | |
| Precuneus, BA 31, Parietal lobule, bleeds into posterior cingulate | Bilateral | 489 | 2 | -72 | 26 | 6.02 | 0.001 |
| | | | -4 | -48 | 32 | | |
| Inferior parietal lobule, Angular | Right | 348 | 46 | -58 | 34 | 5.60 | 0.008 |

| | | | | | | | |
|---|---|---|---|---|---|---|---|
| gyrus, Supramarginal gyrus, Superior temporal gyrus (BA 22) | | | | | | | |
| Lingual gyrus (BA 18), bleeds into Cerebellum anterior lobe | Left | 305 | -18 | -54 | 0 | 5.12 | 0.014 |
| Superior temporal gyrus (BA 39), Supramarginal gyrus, Middle temporal gyrus | Left | 494 | -54 | -62 | 22 | 4.62 | 0.001 |
| Lingual gyrus | Right | 189 | 20 | -52 | 2 | 4.54 | 0.084 (FWE), 0.037 (FDR) |

**<u>Minus C*V2</u>**

| | | | Peak MNI coordinates | | | | |
|---|---|---|---|---|---|---|---|
| **Region** | **Laterality** | **Cluster Size** | **X** | **Y** | **Z** | **Max stat t** | **P Cluster Corrected** |
| Superior parietal lobule, Precuneus / BA 7 | Bilateral | 1921 | 22 | -66 | 58 | 8.46 | <0.001 |
| | | | -6 | -58 | 50 | | |
| Middle occipital gyrus | Right | 833 | 30 | -88 | 6 | 5.63 | <0.001 |
| Postcentral gyrus (BA 2), Inferior parietal lobule (BA 40) | Left | 244 | -46 | -28 | 48 | 5.14 | 0.035 |
| Vermis, Declive, Cerebellum posterior lobe | Bilateral | 331 | 4 | -76 | -28 | 5.00 | 0.01 |
| | | | -2 | -72 | -40 | | |
| Middle occipital gyrus | Left | 341 | -28 | -90 | 6 | 4.60 | 0.008 |

**<u>(C – CntrlC)*V2</u>**

| | | | Peak MNI coordinates Clust thresh = 0.005 | | | | |
|---|---|---|---|---|---|---|---|
| **Region** | **Laterality** | **Cluster Size** | **X** | **Y** | **Z** | **Max stat t** | **P Cluster Corrected** |
| Superior frontal gyrus (BA 9), Anterior cingulate (BA 32) | Bilateral | 496 | 6 | 54 | 32 | 5.91 | 0.090 (FWE), 0.077 (FDR) |
| | | | -6 | 50 | 18 | | |

**(CntrlC – C)*V2**

| | | | Peak MNI coordinates Clust thresh = 0.005 | | | | |
|---|---|---|---|---|---|---|---|
| **Region** | **Laterality** | **Cluster Size** | **X** | **Y** | **Z** | **Max stat t** | **P Cluster Corrected** |
| Cerebellum posterior lobe, Pyramis, Declive, Uvula | Bilateral | 1549 | 6 | -74 | -36 | 4.53 | <0.001 |
| | | | -16 | -62 | -22 | | |

**B*V2**

| | | | Peak MNI coordinates | | | | |
|---|---|---|---|---|---|---|---|
| **Region** | **Laterality** | **Cluster Size** | **X** | **Y** | **Z** | **Max stat t** | **P Cluster Corrected** |
| Inferior parietal lobule | Left | 310 | -50 | -58 | 48 | 4.43 | 0.018 |
| Cindulate gyrus, Posterior cingulate, mid-cingulate | Bilateral | 195 | 6 | -50 | 28 | 4.20 | 0.095 (FWE), 0.078 (FDR) |
| | | | -4 | -30 | 40 | | |

**Minus B*V2**

| | | | Peak MNI coordinates |
|---|---|---|---|

| Region | Laterality | Cluster Size | X | Y | Z | Max stat t | P Cluster Corrected |
|---|---|---|---|---|---|---|---|
| Precuneus / BA 7, Superior parietal lobule | Bilateral | 950 | 8 | -56 | 54 | 5.93 | <0.001 |
| | | | -6 | -66 | 48 | | |
| Middle occipital gyrus | Right | 579 | 34 | -86 | 22 | 4.97 | 0.001 |
| Occipital gyrus | Left | 283 | -24 | -66 | 30 | 4.81 | 0.027 |

**(B – CntrlB)*V2**

No significant clusters.

**(CntrlB – B)*V2**

| | | | Peak MNI coordinates | | | | |
|---|---|---|---|---|---|---|---|
| Region | Laterality | Cluster Size | X | Y | Z | Max stat t | P Cluster Corrected |
| Middle occipital gyrus | Right | 577 | 32 | -86 | 18 | 5.68 | 0.001 |
| Superior parietal lobule, Inferior parietal lobule, BA 7 | Right | 219 | 26 | -56 | 50 | 4.81 | 0.058 (FWE), 0.045 (FDR) |

*Parametric modulation: P1V1*

**(C – B)*P1V1**

| | | | Peak MNI coordinates | | | | |
|---|---|---|---|---|---|---|---|
| Region | Laterality | Cluster Size | X | Y | Z | Max stat t | P Cluster Corrected |
| Lingual gyrus, BA 19 | Right | 271 | 16 | -50 | 4 | 6.59 | 0.022 |

**(B – C)*P1V1**

| | | | Peak MNI coordinates |
|---|---|---|---|

| | | | Clust defining thresh = 0.005 | | | | |
|---|---|---|---|---|---|---|---|
| Region | Laterality | Cluster Size | X | Y | Z | Max stat t | P Cluster Corrected |
| Postcentral gyrus | Left | 614 | -44 | -26 | 56 | 4.71 | 0.028 |

**(C – CntrlC minus B – CntrlB)*P1V1**

No significant clusters.

**(B – CntrlB minus C – CntrlC)*P1V1**

No significant clusters.

**C*P1V1**

| | | | | Peak MNI coordinates | | | |
|---|---|---|---|---|---|---|---|
| Region | Laterality | Cluster Size | X | Y | Z | Max stat t | P Cluster Corrected |
| Superior temporal gyrus, Middle temporal lobe | Right | 368 | 68 | -52 | 12 | 6.07 | 0.006 |
| Lingual/ Parahippocampal gyrus/ BA 30 | Right | 279 | 20 | -52 | 2 | 4.88 | 0.02 |
| Medial frontal gyrus / BA 10 (Somewhat bleeds into anterior cingulate) | Left | 259 | -6 | 50 | 4 | 4.79 | 0.027 |
| Lingual gyrus | | 163 | -20 | -68 | -8 | 4.70 | 0.128 (FWE), 0.079 (FDR) |

**Minus C*P1V1**

| | | | | Peak MNI coordinates | | | |
|---|---|---|---|---|---|---|---|
| Region | Laterality | Cluster Size | X | Y | Z | Max stat t | P Cluster Corrected |

| | | | | | | | |
|---|---|---|---|---|---|---|---|
| Precuneus, Superior parietal lobule / BA 7 | Bilateral | 1286 | 8 | -58 | 58 | 6.79 | <0.001 |
| | | | -6 | -60 | 52 | 6.71 | |
| Postcentral gyrus / BA 2, Precentral gyrus / BA 4 | Left | 343 | -50 | -32 | 56 | 4.85 | 0.008 |
| Middle occipital gyrus | Right | 266 | 24 | -94 | 8 | 4.59 | 0.025 |
| Inferior parietal lobule / BA 40 | Right | 248 | 44 | -32 | 42 | 4.49 | 0.032 |

Additional clusters with lower clust defining thresh.

| | | | **Peak MNI coordinates Clust def thresh = 0.005** | | | | |
|---|---|---|---|---|---|---|---|
| **Region** | **Laterality** | **Cluster Size** | **X** | **Y** | **Z** | **Max stat t** | **P Cluster Corrected** |
| Precentral gyrus | Right | 519 | 32 | -14 | 60 | 4.12 | 0.062 (FWE), 0.015 (FDR) |
| Cerebellum_6, Cerebellum anterior lobe | Left | 300 | -26 | -48 | -28 | 4.03 | 0.345 (FWE), 0.066 (FDR) |

**(C – CntrlC)*P1V1**

No significant clusters.

**(CntrlC – C)*P1V1**

| | | | **Peak MNI coordinates Clust def thresh = 0.005** | | | | |
|---|---|---|---|---|---|---|---|
| **Region** | **Laterality** | **Cluster Size** | **X** | **Y** | **Z** | **Max stat t** | **P Cluster Corrected** |
| Vermis, Cerebellum posterior lobe | Right | 1243 | 2 | -74 | -34 | 4.27 | 0.001 |

| | | | | | | | |
|---|---|---|---|---|---|---|---|
| Thalamus, Lentiform nucleus, Putamen | Right | 479 | 22 | -16 | 16 | 4.21 | 0.103 (FWE), 0.066 (FDR) |
| Cerebellum_8, Cerebellum posterior lobe | Left | 404 | -24 | -66 | -54 | 4.14 | 0.179 (FWE), 0.066 (FDR) |

**B*P1V1**

| | | | Peak MNI coordinates (Clust def thresh = 0.008) | | | | |
|---|---|---|---|---|---|---|---|
| **Region** | **Laterality** | **Cluster Size** | **X** | **Y** | **Z** | **Max stat t** | **P Cluster Corrected** |
| Middle frontal gyrus / BA 10 / Middle orbital frontal gyrus, Anterior cingulum | Bilateral | 1053 | -10 | 40 | -8 | 4.53 | 0.012 |
| | | | 14 | 42 | 4 | | |

**Minus B*P1V1**

| | | | Peak MNI coordinates Clust def thresh = 0.005 | | | | |
|---|---|---|---|---|---|---|---|
| **Region** | **Laterality** | **Cluster Size** | **X** | **Y** | **Z** | **Max stat t** | **P Cluster Corrected** |
| Precuneus, Superior parietal lobule | Right | 1748 | 8 | -56 | 54 | 5.61 | <0.001 |

**(B – CntrlB)*P1V1**

No significant clusters.

**(CntrlB – B)*P1V1**

No significant clusters.

*Parametric modulation: P3V3*

**(C – B)*P3V3**

| | | | Peak MNI coordinates Clust def thresh = 0.008 | | | | |
|---|---|---|---|---|---|---|---|
| **Region** | **Laterality** | **Cluster Size** | **X** | **Y** | **Z** | **Max stat t** | **P Cluster Corrected** |
| Inferior temporal gyrus | Left | 806 | -44 | -24 | -20 | 4.38 | 0.034 |

(B – C)*P3V3

No significant clusters.

**(C – CntrlC minus B – CntrlB)*P3V3**

| | | | Peak MNI coordinates Clust def thresh = 0.008 | | | | |
|---|---|---|---|---|---|---|---|
| **Region** | **Laterality** | **Cluster Size** | **X** | **Y** | **Z** | **Max stat t** | **P Cluster Corrected** |
| Thalamus, Lentiform Nucleus, Globus Palladus | Left | 636 | -10 | -4 | -2 | 5.71 | 0.098 |

**(B – CntrlB minus C – CntrlC)*P3V3**

No significant clusters.

**C*P3V3**

| | | | Peak MNI coordinates Clust def thresh = 0.005 |
|---|---|---|---|

| Region | Laterality | Cluster Size | X | Y | Z | Max stat t | P Cluster Corrected |
|---|---|---|---|---|---|---|---|
| Inferior frontal gyrus | Left | 779 | -54 | 14 | 26 | 5.32 | 0.014 |

**Minus C*P3V3**

No significant clusters.

**B*P3V3**

No significant clusters.

**Minus B*P3V3**

No significant clusters.

**(C-CntrlC)*P3V3**

No significant clusters.

**(CntrlC-C)*P3V3**

No significant clusters.

**(B-CntrlB)*P3V3**

No significant clusters.

**(CntrlB-B)*P3V3**

No significant clusters.

*Parametric modulation: Value of gamble according to policies 2 and 5.*

**C*Pol5 – B*Pol2**

No significant clusters.

**B*Pol2 – C*Pol5**

| | | | Peak MNI coordinates | | | | |
|---|---|---|---|---|---|---|---|
| **Region** | **Laterality** | **Cluster Size** | **X** | **Y** | **Z** | **Max stat t** | **P Cluster Corrected** |
| Cerebellum anterior lobe, Culmen | Left | 230 | -26 | -46 | -26 | 6.05 | 0.055 (FWE), 0.072 (FDR) |
| Postcentral gyrus (BA 2, BA 3), Precentral gyrus | Left | 741 | -48 | -28 | 56 | 5.54 | <0.001 |

### Supplementary Discussion

#### 1. Consistency as a measure of commitment to a single policy

Interpreting consistency measure as indexing commitment to a single policy throughout the task, at the level of individual participants, is supported by the observation that participants who reported higher difficulty (Choice – Best) in making decisions, also showed lower consistency. Higher difficulty suggests greater conflict between options, that is, multiple policies in this case and hence, may explain its negative relation with lower consistency. The interpretation aligns with smaller relative RT being associated with a greater relative consistency (Figure 4).

#### 2. Task order effects on the consistency measure

Participants performed two blocks of the same task first, followed by two blocks of the other later (order 1: Choice (x 2) → Best (x 2); order 2: reverse of order 1). The effect of task order is not surprising as after the first task, participants would become more familiar with the stimuli and the general demands of a decision-making task, making it easier to converge on a policy. This may also be facilitated by fatigue. The consistency in Best when it is conducted first was not different from when it is conducted the second. It was only the consistency in Choice that is affected by order such that it shows greater consistency when it is conducted second, after the Best, than when it is conducted first. This suggests that the 'objective' decision-making mode (Best task) might be 'sticky' and interfered with the performance in Choice, such that it led to a quicker convergence to single policy in the Choice blocks, when they were conducted after Best.

#### 3. Other objectivity measures: Distortion and non-randomness

Distortion measure quantified the extent to which externally provided (or the 'objective') information, like the values and probabilities associated with the options, was distorted. Non-randomness measure quantified the extent to which the decision in a trial was predicted with certainty by a policy. These were measures of objectivity, in addition to the consistency measure, and were obtained at the level of individual participants. These did not show significant differences between the Choice and the Best conditions. As noted in main article, consistency measures showed order effects (Figure 2H) which could be attributed to a greater familiarity or fatigue. Neither non-randomness and distortion measure nor the measures of alignment with optimal policies showed any order effects, suggesting that, in terms of these measures, increased familiarity with the task or fatigue did not significantly affect what they measured.

### Appendix

#### Task-strategy questionnaire

{Text in curly brackets was not shown to the participants. Participants completed this questionnaire in a word document on a computer. They were free to use as much space as they wanted to write their responses.}

1. Please describe your thought process and the strategy you used in responding to trials where you were asked to indicate the best option. Please feel free to write as much or as little as you deem necessary to describe your thought process and strategy. The space provided below does not reflect our expectations regarding the length of the description. You may use the back of the page if you need more space to write.

2. Please describe your thought process and the strategy you used in responding to trials where you were asked to choose an option. Please feel free to write as much or as little as you deem necessary to describe your thought process and strategy. The space provided below does not reflect our expectations regarding the length of the description. You may use the back of the page if you need more space to write.

3. Was there any difference in your thought process or strategy when you were asked to indicate the best option vs when you were asked to choose? If so, please describe. Please feel free to write as much or as little as you deem necessary to describe your thought process and strategy. The space provided below does not reflect our expectations regarding the length of the description. You may use the back of the page if you need more space to write.

4. Please describe your thought process or strategy when you were provided an instruction to indicate an option as best or choose a specific option. Did you think about only the option mentioned in the instruction or did you also think about and considered selecting the other option? Were there any differences in your experience of or in your approach to the task when you were instructed to indicate an option as best or when you were instructed to choose an option?

5. Please provide a rating on a scale of 1-10 for each of the three tasks on the following grounds:

| Task | Amount of attention or focus required to complete the task<br>1 – Minimal focus<br>10 – All the focus on the task | Difficulty level of the decision-making process<br>1 – Did not even feel like a decision, Low effort<br>10 – Very involved decision; High effort |
|---|---|---|
| Task where you indicated the BEST option | | |
| Task where you CHOSE an option | | |
| Task where you indicated the **best** option as INSTRUCTED | | |
| Task where you **chose** as INSTRUCTED | | |

6. Expected value: Average outcome of a probabilistic option. For example, consider a gamble that leads to an outcome of 100 points with 20% chance, 60 points with 50% chance, and -50 with chance 30% as shown below. Its expected value is 35.

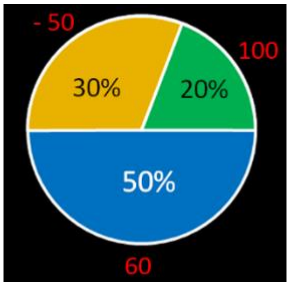

Expected value of the gamble = (100 x (20/100)) + (60 x (50/100)) + ((-50) x (30/100)) = (100 x 0.2) + (60 x 0.5) + ((-50) x 0.3) = 20 + 30 - 15 = 35

Please indicate if you were familiar with the concept of expected value when you performed the task? Please indicate the level of familiarity on a scale of 1-10. Here, 1 – Completely unaware of this concept; 10 – Aware of the concept and comfortable with calculating the expected value.

1 2 3 4 5 6 7 8 9 10

7. Please indicate if you thought about the following factors in the trials where you indicated the best option and how important were those factors to you on a scale of 1-10. Please mark

zero for factors you did not think about. Please mark a number between 1 and 10 indicating their importance to you for the factors that you thought about while deciding.

0 – Did not think about the factor
1 – Not at all important or barely affected my decision
10 – Extremely important or played a critical role in my decision

| Factor | Trials where you indicated the **best** option |
|---|---|
| Probability of the negative outcome.<br>Example: 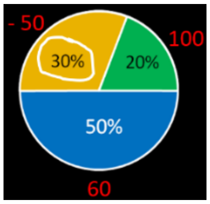  | 0 1 2 3 4 5 6 7 8 9 10 |
| Value of the negative outcome.<br>Example: 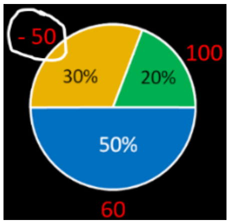  | 0 1 2 3 4 5 6 7 8 9 10 |
| Probability of the highest possible outcome of the gamble.<br>Example: 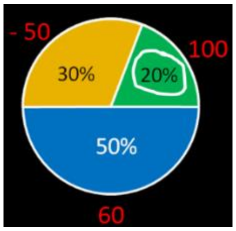  | 0 1 2 3 4 5 6 7 8 9 10 |
| Value of the highest possible outcome of the gamble. | 0 1 2 3 4 5 6 7 8 9 10 |

| | |
|---|---|
| Example: 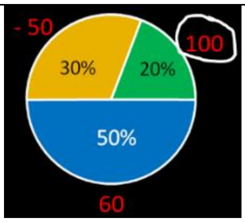 | |
| Probabilities associated with positive outcomes of the gambles taken together.<br>Example: 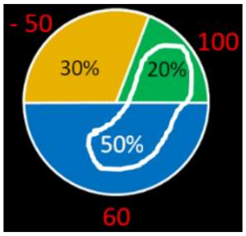 | 0 1 2 3 4 5 6 7 8 9 10 |
| Value of positive outcomes of the gamble taken together.<br>Example: 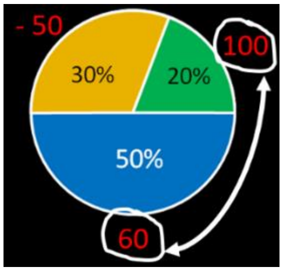 | 0 1 2 3 4 5 6 7 8 9 10 |
| Expected value of the gamble. | 0 1 2 3 4 5 6 7 8 9 10 |

8. Please indicate if you thought about the following factors in the trials where you were asked to choose an option and how important were those factors to you on a scale of 1-10. Please mark zero for factors you did not think about. Please mark a number between 1 and 10 indicating their importance to you for the factors that you thought about while deciding.

   0 – Did not think about the factor
   1 – Not at all important or barely affected my decision

10 – Extremely important or played a critical role in my decision

| Factor | Trials where you **chose** an option |
|---|---|
| Probability of the negative outcome.<br>Example: 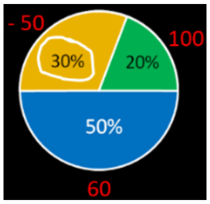  | 0 1 2 3 4 5 6 7 8 9 10 |
| Value of the negative outcome.<br>Example: 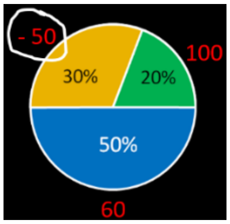  | 0 1 2 3 4 5 6 7 8 9 10 |
| Probability of the highest possible outcome of the gamble.<br>Example: 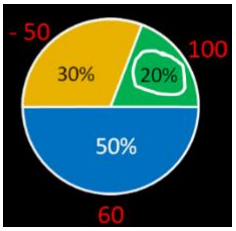  | 0 1 2 3 4 5 6 7 8 9 10 |
| Value of the highest possible outcome of the gamble.<br>Example: 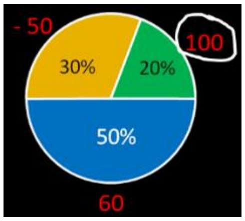  | 0 1 2 3 4 5 6 7 8 9 10 |

| | |
|---|---|
| Probabilities associated with positive outcomes of the gambles taken together.<br><br>Example: 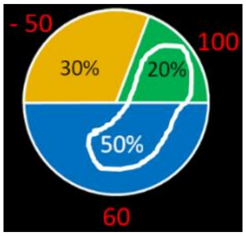 | 0 1 2 3 4 5 6 7 8 9 10 |
| Value of positive outcomes of the gamble taken together.<br><br>Example: 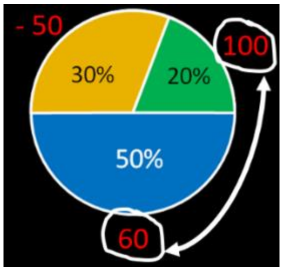 | 0 1 2 3 4 5 6 7 8 9 10 |
| Expected value of the gamble. | 0 1 2 3 4 5 6 7 8 9 10 |

9. Please indicate whether you agree or disagree with the following statements about how you performed the task, using the following scale:

| Strongly disagree | Disagree | Neither agree nor disagree | Agree | Strongly Agree |
|---|---|---|---|---|
| 1 | 2 | 3 | 4 | 5 |

________ In trials where I was asked to indicate which option is <u>best</u>, I thought about the options before making my choice, instead of randomly making a choice without thinking.

_______ In trials where I was asked to indicate which option is <u>best</u>, my answers were based on my previous experience with the options instead of information about the options shown on the screen.

_______ In trials where I was asked to choose an option, I thought about the options before indicating my choice, instead of randomly making a choice without thinking.

______ In trials where I was asked to choose an option, my answers were based on my previous experience with the options instead of information about the options shown on the screen.

______ I used the same strategy in both the trials where I was asked to indicate an option that was best and the trials where I was asked to choose an option.

______ I used different strategies in the trials where I was asked to indicate the option that is best and the trials where I was asked to choose an option.

______ It was hard to distinguish between trials where I was asked to choose an option and the trials where I was asked to indicate which option is best.

______ I was tracking the total points earned in trials where I was asked to indicate which option is best.

______ I was tracking the total points earned in trials where I was asked to choose an option.

10. Please indicate the frequency of the following conditions as you worked through the task, using the following scale:

| | | | | | | |
|---|---|---|---|---|---|---|
| 90 – 100% | 80 – 90% | 70 – 80% | …………….. | 20 – 30% | 10 – 20% | 0 – 10% |
| 10 | 9 | 8 | …………….. | 3 | 2 | 1 |

_______ In trials where I was asked to choose an option, I chose the option that I thought was objectively the best.

_______ In trials where I was asked to choose an option, I chose the option that I did not necessarily think was objectively the best.

_______ In trials where I was asked to choose an option, I chose the option that I did not necessarily prefer but the option seemed objectively best for my reward total.

_______ In trial where I was asked to choose an option, I did not think about which option was objectively the best.

#### Mathematics exam

{Text in curly brackets was not shown to the participants. The test was conducted in Qualtrics and the time taken to respond to each question was also noted.}

Please answer the following questions. The time taken to finish each question will be noted.

{Approximation}

1. Please indicate which option is closest to 48*0.5
   A. 10
   B. 18
   C. 35
   D. 25

2. Please indicate which option is closest to 102*0.3
   A. 16
   B. 30
   C. 40
   D. 56

3. Please indicate which option is closest to 500*0.19
   A. 50
   B. 125
   C. 75
   D. 100

4. Please indicate which option is closest to 20*0.082
   A. 0.8
   B. 1
   C. 1.6
   D. 2.4

Please answer the following questions. You will have 5 seconds to complete each question.

{Probability}

5. Imagine you bought a lottery ticket to win $1000. Your probability of winning is 0.3. What is the probability that you will not win? __________

6. Imagine that there are 8 people in a neighborhood. The chances that any of them is a zombie are 50%. How many people out of 8 would you expect to be zombies? __________

7. Imagine you bought a lottery ticket. It is a different type of lottery such that you may win a food processor with probability 0.2 or you may win a vacuum cleaner with probability 0.1 or you may win nothing with probability 0.7. What is the probability of winning something from the lottery? _________

In each of the following, find the expected values (X) of the gambles. Here X = sum of product of probabilities and values.

Example:

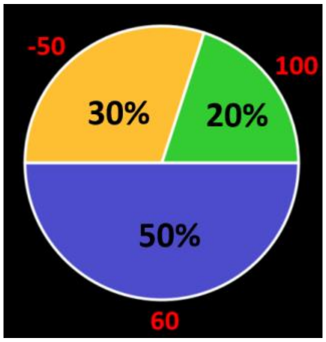

X for this gamble is, $X = (60 * 0.5) + (100 * 0.2) + ((-50) * 0.3)$

$$= (30) + (20) + (-15)$$

$$= 35$$

8. Find X.

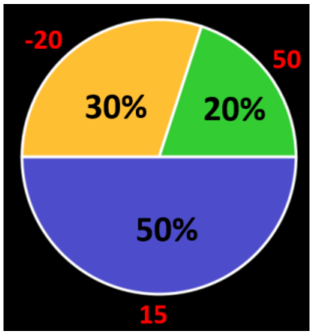

Your answer: ______________

9. Find X.

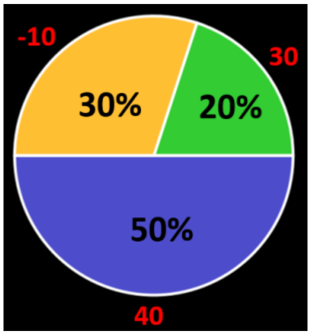

Your answer: _____________

10. Find X.

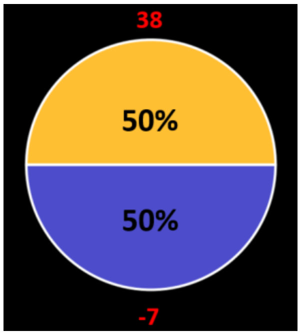

Your answer: _____________

11. Find X

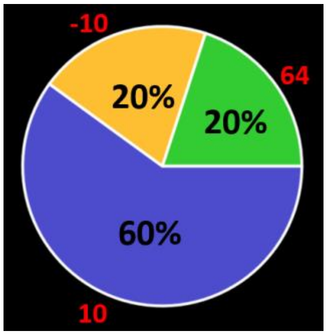

Your answer: _____________

12. Find X.

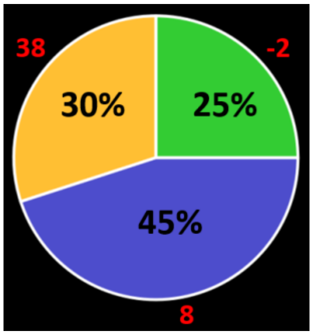

Your answer: _____________

{Correct answers:

1. **D**, 2. **B**, 3. **D**, 4. **C**, 5. **0.7**, 6. **4**, 7. **0.3**, 8. **11.5**, 9. **23**, 10. **15.5**, 11. **16.8**, 12. **14.5**}
